# FAM83H regulates postnatal T cell development through thymic stroma organization

**DOI:** 10.1101/2025.05.26.656197

**Authors:** BM Ogan, V Forstlová, L Dowling, M Šímová, K Vičíková, V Novosadová, F Špoutil, S Červenková, M Procházková, J Turečková, O Fedosieieva, J Lábaj, P Nickl, K Křížová, J Procházka, R Sedláček, J Balounová

## Abstract

Family of Sequence Similarity 83H (FAM83H/ SACK1H) is primarily expressed in epithelial cells, where it interacts with casein kinase 1 (CK1) and keratins to regulate cytoskeletal organization, cell proliferation, and vesicular trafficking. Mutations in *FAM83H* are known to cause amelogenesis imperfecta, highlighting its critical role in enamel formation. We generated *Fam83h*-deficient mice (*Fam83h^em2(IMPC)Ccpcz^, Fam83h^-/-^*) and mice lacking a part of the N-terminal CK1-binding domain (*Fam83h*^Δ*87/*Δ*87*^). Consistent with other *Fam83h*-deficient models, these mice are subviable, smaller in size, and exhibit a sparse, scruffy coat, scaly skin, general weakness, and hypoactivity. Notably, both strains show impaired lymphoid cell development in early postnatal life. In the thymus, *Fam83h* expression is confined to thymic epithelial cells (TECs), and its deficiency in stromal cells results in disrupted thymic architecture and severe block in the expansion of DN3 (double-negative stage 3) T cells, ultimately leading to insufficient T cell production. Single-cell transcriptomic analysis reveals that *Fam83h^-/-^* cortical TECs (cTECs) express reduced levels of the TEC master regulator *Foxn1*, and its multiple downstream target genes, suggesting a critical role for FAM83H likely in coordination with CK1in cTEC maturation.

**Highlights:**

*Fam83h*-deficient mice exhibit multiple epithelial defects, but no obvious enamel defects
*Fam83h* deficiency disrupts lymphocyte development
*Fam83h* deficiency impairs thymic epithelial cell maturation and thymus function

## 1. Introduction

The Family with Sequence Similarity 83 Member H (FAM83H, or SACK1H) plays a crucial role in enamel formation by maintaining the structural integrity of ameloblasts (1, 2). Truncating mutations in *FAM83H* gene disrupt ameloblast function, leading to defective enamel mineralization and the clinical manifestation of autosomal dominant hypocalcified amelogenesis imperfecta (ADCAI). These mutations, often nonsense or frameshift variants, result in truncated proteins that are likely to disrupt the functional C-terminus (3, 4).

The FAM83 protein family, comprising eight members, is evolutionarily conserved across vertebrates yet remains relatively uncharacterized. All members share a conserved N-terminal casein kinase 1 (CK1) binding domain, while their unique C-terminal regions vary in length and sequence (5). FAM83 proteins thus may bind and sequester CK1 isoforms at discrete subcellular locations, either directing CK1 toward or away from substrates, or serving as a physical bridge or barrier between CK1 and its targets (6)..FAM83H interacts with CK1α at its N-terminus and with keratins at its C-terminus, playing a key role in organizing keratin filaments by recruiting CK1α (7). Thus, FAM83H supports mechanical stability of epithelial cells, as well as epithelial cell polarity through its effects on keratin organization and desmosome assembly (1, 7). Beyond ameloblasts, FAM83H is predominantly expressed in other epithelial cells, including those of the intestine, skin, and mammary gland, but appears to be absent in immune cells (8, 9). Additionally, FAM83H expression correlates with poor prognosis various cancers (10), including squamous cell carcinoma (11), gastric carcinoma (12), and breast cancer (13). Beyond enamel defects, *Fam83h* deficiency in mice leads to broader physiological abnormalities. For example, Zheng *et al*. demonstrated that a knock-in mouse carrying the *Fam83h* c.1186C>T (p.Q396*) mutation - analogous to the human ADCAI *FAM83H* c.1192C>T variant developed forelimb swelling and inflammation (2).

Lymphocyte development, particularly T cell differentiation, begins with hematopoietic stem cells in the bone marrow that generate lymphoid progenitors (14). T cell precursors migrate to the thymus, a primarily epithelial organ, where they mature (15). The thymus is dynamically remodeled by incoming hematopoietic progenitors (16). Reciprocal interactions between hematopoietic and stromal cells (known as lymphoepithelial crosstalk) support the stromal cell maturation in two main types—cortical and medullary thymic epithelial cells (cTECs/ mTECs) which then provide signals needed for T cell development (17). Specifically, cTECs coordinate the early stages of T cell development and positive selection (18), and mTECs mediate negative selection (15). Recent single-cell RNA-sequencing (scRNA-seq) studies have revealed unexpected TEC heterogeneity and provided new insights into their development and function under homeostatic conditions (19–21). In these studies, FAM83H mRNA was found to be expressed in TEC, especially cTECs and intertypical TECs (20, 21). Multiple molecular pathways govern thymic development, with the Wnt/β-catenin signaling pathway increasingly recognized as essential (22). Notably, overexpression of β-catenin overexpression in TECs reduces thymic cellularity and TEC numbers, highlighting the importance of finely tuned Wnt signaling (23).

FAM83H has emerged as a potential regulator of the Wnt/β-catenin pathway (24). Kuga *et al.* reported that *Fam83h* deletion led to reduced Wnt/β-catenin activity, evidenced by decreased expression levels of CK1α, CK1ε, and β-catenin (25).

To date, the role of FAM83H in the immune system has not been studied. We demonstrate that FAM83H expression in stromal cells significantly affects lymphocyte production. In the thymus, *Fam83h* deficiency disrupts stromal cell architecture, impairs DN (double-negative) T cell expansion in the cortex, and results in severely reduced T cell output. The same outcome is observed in animals with a truncated form of FAM83H lacking part of the CK1-binding domain, suggesting a critical role of the Fam83h-CK1α interaction in this process. Additionally, *Fam83h*-deficient cTECs show elevated expression of circadian rhythm-related genes, and reduced expression of the TEC master regulator *Foxn1* and its multiple downstream targets, suggesting a broader impact on TEC maturation and function.

In summary, FAM83H, in coordination with CK1, is essential for organizing the keratin cytoskeleton in TECs. This structural role together with the impact on TEC maturation supports proper thymic architecture and is critical for sustaining normal T cell development.

## 2. Results

### 2.1 *Fam83h* is expressed in epithelial cells across multiple tissues

*Fam83h* is enriched in epithelial cells of the gastrointestinal tract, skin, bladder, tongue, pancreas, and mammary gland **(Figure 1A)** (26). We detected *Fam83h* mRNA expression in multiple organs, with the intestine showing unexpectedly higher levels than ameloblasts. Additionally, low levels of *Fam83h* mRNA were observed in the spleen and bone marrow (BM) **(Figure 1B).**

**Figure 1.**
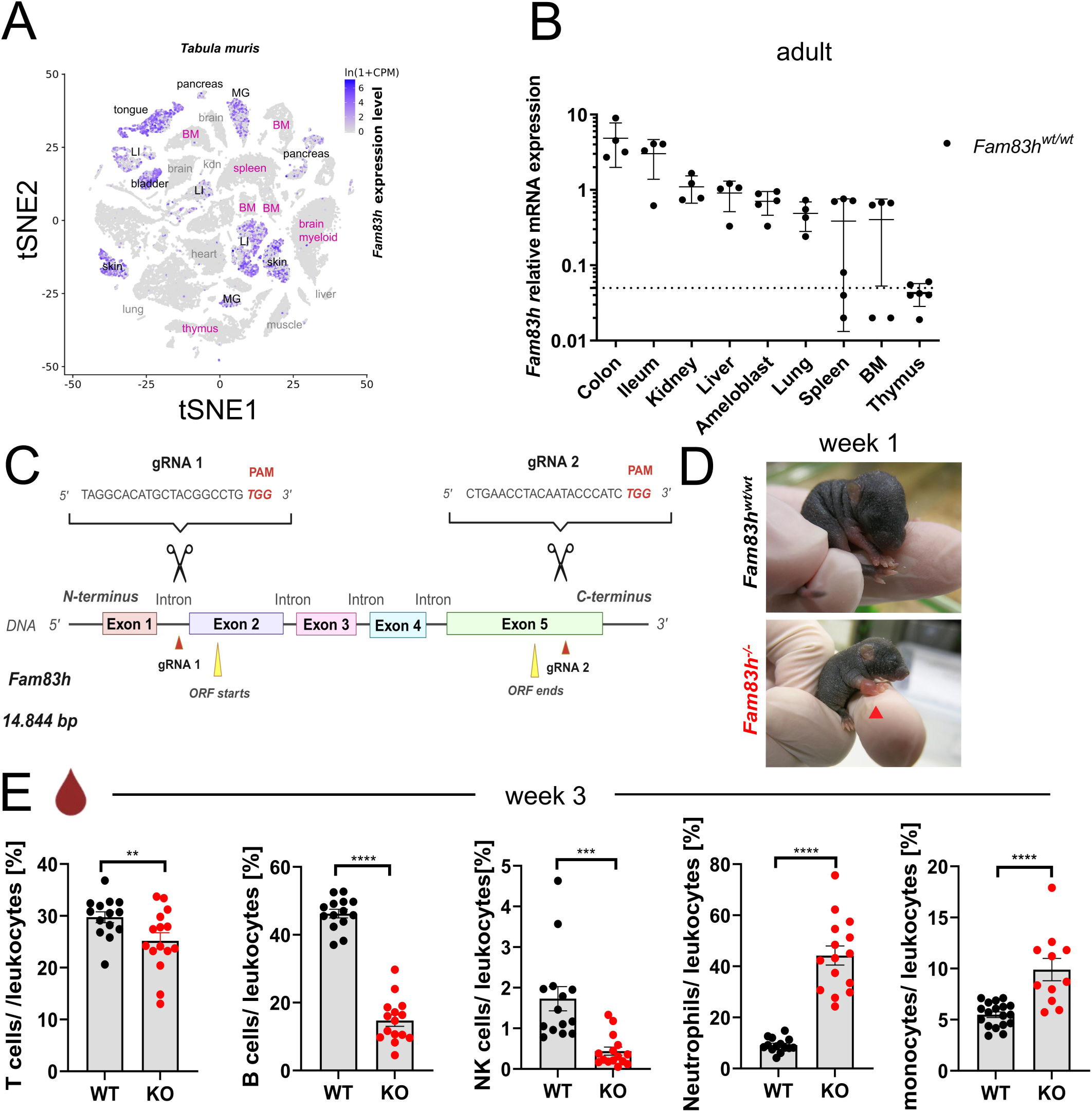
*Fam83h* deficiency leads to dysregulation of early postnatal hematopoiesis and growth defects. (A) Expression of *Fam83h* in mouse tissues (Tabula muris, https://www.czbiohub.org/sf/tabula-muris/). (B) Relative mRNA expression levels of *Fam83h* in selected tissues from *Fam83h^wt/wt^*mice, normalized to the reference gene *Casc3* (RT-qPCR). Symbols indicate individual mice. Data are presented as mean ± SD, *n*=4 in GraphPad (version 9). (C) Generation of *Fam83h^-/-^* mice – Schematic representation of the CRISPR/Cas9-mediated knockout strategy for *Fam83h*, targeting regions upstream of exon 2 and within exon 5 using two sgRNAs is depicted (created by BioRender). (D) By postnatal week 1, *Fam83h^-/-^* mice exhibit forepaw swelling compared to *Fam83h^wt/wt^* littermates. (E) At postnatal week 3, *Fam83h^-/-^* mice show altered hematopoietic profiles in peripheral blood, assessed by flow cytometry (FCM). The gating strategy is depicted in Figure S8. The graph represents 6 independent experiments. Symbols denote individual mice, with bars indicating mean ± SEM, unpaired *t-test*, p-values: **P ≤ 0.01, ****P ≤ 0.0001.

### 2.2. *Fam83h*-deficient mice display a spectrum of epithelia-related phenotypic abnormalities

We generated *Fam83h*-deficient mice (*Fam83h^em2(IMPC)Ccpcz^,* hereafter *Fam83h^-/-^*) and mice lacking part of the N-terminal casein kinase 1 (CK1)-binding domain (*Fam83h*^Δ*87/*Δ*87*^) **(Figures 1C, and S7A)**. Consistent with other *Fam83h*-deficient models, *both Fam83h-*deficient strains are subviable, smaller in size, and display a sparse, scruffy coat, scaly skin, generalized weakness, hypoactivity, and develop skin and bone lesions early in life **(Figures 1D, S1A, B, and D)** (2, 27, 28). Notably, these manifestations appeared early in life, with few animals surviving beyond 2-4 weeks of age **(Figure S1A, right panel, and S7B)**. Histological analysis revealed forelimb edema (**Figure 1D**) with immune cell infiltrates (**Figure S1B**). Although dental abnormalities were not observed, bone mineral density appeared reduced in *Fam83h^-/-^*mice based on pseudo color analysis of micro computed tomography (µCT) images (**Figure S1B, C**).

### 2.3 FAM83H modulates hematopoietic cell output

Unexpectedly, *Fam83h-*deficient *-* mice exhibited increased myeloid and decreased lymphoid cells in peripheral blood during early postnatal life, (week 3) **(Figures 1E, S1E, and S7C)**. In animals surviving beyond week 3, this imbalance began to normalize by week 8 and returned to baseline by week 12 (**Figure S1E)**. Flow cytometry (FCM) analysis of *Fam83h^-/-^* BM progenitors (**Figures S2-3**) conducted during weeks 1-3, showed that lymphoid progenitor production, reduced already at week 1, became even more impaired by week 3 **(Figures S2).** Specifically, common lymphoid progenitors (CLPs), which sit at the top of the lymphoid hierarchy, were significantly reduced in the BM of *Fam83h^-/-^* animals at week 3. This decline propagated downstream, affecting B and NK cell progeny, with pre-BII cell fraction reduced by more than 1,000-fold. **(Figure S2A-C)**. In absolute numbers, we observed a highly significant decrease in pre-BII cells, pro-B cells, immature B cells, and mature NK cells already at week 2 (**Figures S2D, E**). At week 2, the reduction in lymphoid progenitors was reflected by a reduced total BM cell count (**Figure S3A**). In contrast, erythro-myeloid progenitors remained largely unaffected (**Figures S3A, B**). Although the proportions of neutrophils and monocytes were increased, their absolute numbers were not significantly altered (**Figure S3B**). Among hematopoietic stem cells and progenitors (HSPCs), absolute counts of phenotypical LT-LSK cells were decreased in *Fam83h^-/-^* BM at week 2 (**Figure S3C**). Figure S3D illustrates the frequencies of individual BM hematopoietic cell populations at week 3.

### 2.4 *Fam83h* deficiency disrupts thymic architecture and impairs T cell production

The deficiency in lymphoid progenitors was even more pronounced in the thymus. Starting at week 2, *Fam83h^-/-^* mice exhibited significantly reduced relative thymus-to-body weight ratio compared to wild-type (wt) controls **(Figure 2A, left panel and S7D)**. Additionally, absolute thymocyte counts were decreased in *Fam83h*-deficient animals from week 2 (**Figure 2A, right panel, Figure S7D, right panel**). Reduced proliferation was evident in both TECs and thymocytes **(Figure 2B)**.

**Figure 2.**
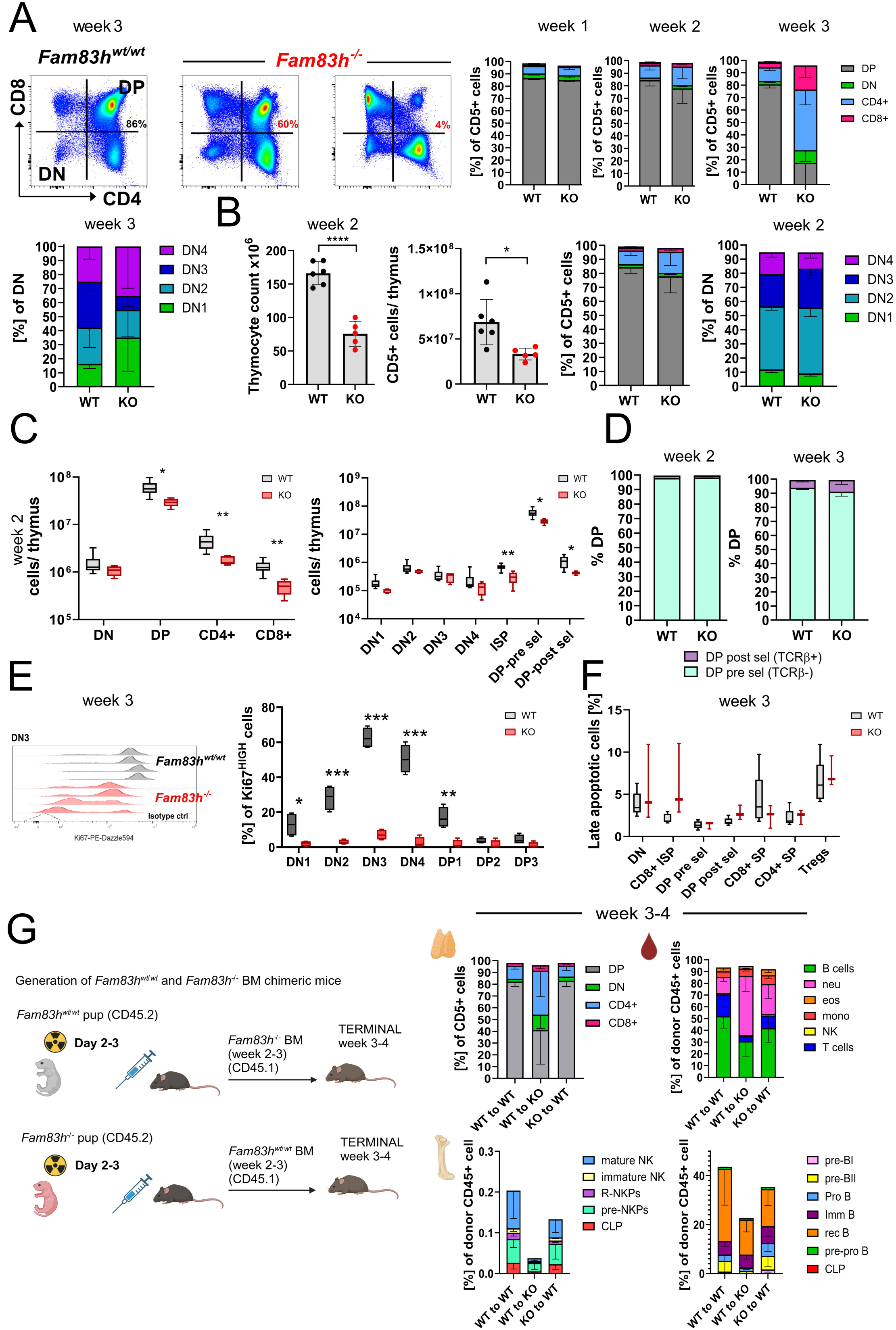
*Fam83h* deficiency leads to altered thymic architecture in early-postnatal thymus. (A) Relative thymus-to-body weight ratio (left) and total thymic cellularity (right) were assessed at weeks 1, 2, and 3. The graph represents 3 independent experiments for week 1, 11 for week 2, and 6 for week 3. n=8-16 for *Fam83h^wt/wt^* and 5-16 for *Fam83h^-/-^*. Symbols indicate individual mice, mean ± SD, unpaired *t-test.* p-values: ns (P > 0.05), ***P ≤ 0.001, ****P ≤ 0.0001. (B) The proportions of proliferating, Ki67HIGH cells within the TEC and thymic CD45⁺ populations were measured by FCM (for gating hierarchy see Figure S19 at week 2–3 of age, *n*=2 for *Fam83h^wt/wt^* and *n*=4 for *Fam83h^-/-^*, unpaired *t-test* (box and whiskers, median/ min to max), p-values: **p ≤ 0.01. (C) Hematoxylin and Eosin (H&E) staining of old *Fam83h^wt/wt^* and *Fam83h^-/-^*thymi. . Imaged by Leica DM3000 LED microscopes at 20x/40x magnification. Representative images from 3 independent experiments. Scale bar = 1mm, 500 μm and 200 μm (D) Double immunofluorescence staining of *Fam83h^wt/wt^ and Fam83h^-/-^* thymi for keratin-8 (cTEC, red) and keratin-5 (mTEC, green) enables distinguishing cortical (red) and medullary (green) areas in 2-3 week old thymi. Imaged by ZEISS Axio Imager Z.2. DAPI staining marks nuclei (blue), aiding in cortex-medulla distinction. Representative images from 3 independent experiments. Scale bar = 100 μm and 10 μm Total thymic and cortical area was quantified from sections. Bars represent mean ± SD, n=9, unpaired *t-test.* p-values: ***P ≤ 0.001. Percentage of cTECs and mTECs from EpCAM+ cells was determined by FCM. The gating strategy is shown in Figure S17. Bars represent mean ± SD, n= 8, unpaired *t-test,* p-values: **P ≤ 0.01. (E) Cryosections from week 3 old *Fam83h^wt/wt^ and Fam83h^-/-^* thymi were stained for E-cadherin (red) and keratin-5 (mTEC, green), analyzed using by spinning disk confocal microscope (SDCM). Nikon CSU-W1. Scale bar = 10 μm. (F) Cryosections from week 3 old *Fam83h^wt/wt^ and Fam83h^-/-^* thymi were stained for ZO-1 (red) and E-cadherin (green), imaged with SDCM Nikon CSU-W1. Scale bar = 10 μm.

Histological hematoxylin-eosin and immunofluorescence (IF) analyses revealed severely disrupted thymic architecture in *Fam83h^-/-^*animals. The cortical region was markedly thinner, and medullary regions were poorly defined **(Figures 2C, D)**. Image quantification analysis showed that the cortical area was reduced in *Fam83h^-/-^*animals at week 2. However, FCM analysis revealed higher frequency of cTECs among EpCAM^+^ cells in *Fam83h^-/-^* thymi (**Figure 2D, right panels**). Thus, the reduced cortical area observed in the absence of FAM83H is likely given by lack of developing thymocytes, rather than TECs.

TECs appeared more rounded, with fewer protrusions and an increased co-expression of keratin 5 and keratin 8, indicating impaired TEC differentiation and organization **(Figures 2D-H, and S4A, B)**. Additionally, E-cadherin and zonulin staining revealed less organized stroma in the absence of FAM83H (**Figures 2F, H**).

### 2.5 *Fam83h* deficiency impairs proliferation of DN T cell progenitors in the thymic cortex

FCM analysis of thymocytes at week 3 revealed a dramatic reduction in the double-positive (DP, CD4^+^ CD8^+^) thymocyte population in *Fam83h^-/-^* mice **(Figure 3A, S7E)**. While some mice retained near-normal DP frequencies, others showed a severe loss, with DP cells comprising as little as 4% compared to ∼86% in *Fam83h^wt/wt^* controls **(Figure 3A)**. Among all subsets, DP cells were the most affected. The frequencies of DN1-4 subset were even more variable among *Fam83h^-/-^* individuals, with a decrease in DN3 subset (**Figure 3A).** In absolute counts, *Fam83h*-deficient thymi contained half as many cells as their *Fam83h^wt/wt^*littermates already by week 2 (**Figure 3B)**. Notably, the proportions of T cell developmental stages were only minimally altered at this age (**Figure 3C, left panel)**. The largest cellularity reduction was observed in immature single-positive (ISP) CD8 thymocytes and of both pre- and post-selection DP thymocytes (**Figure 3C, right panel**). However, the percentages of pre-selection and post-selection (TCRβc) DP cells at week 2 and week 3 remain unchanged (**Figure 3D**). These data indicate that β-selection and positive selection are not intrinsically impaired in *Fam83h*-deficient mice. Finaly, Ki67 staining confirmed a significant decrease in proliferation among double-negative (DN) thymocytes, particularly in DN3–DN4 stages, which normally undergo extensive expansion (**Figures 3E and S4C**). Apoptosis rates, however, were not significantly altered across progenitor subsets **(Figure 3F).** While thymic Tregs were either not significantly altered, or slightly decreased, peripheral blood Treg fraction was increased and among all blood T cell subsets, resting phenotype prevailed (**Figures S4D, E**). The altered frequencies of developing T cells are depicted in Figure S4F. To determine whether lymphopenia originated from *Fam83h* deficient stromal cells or hematopoietic cells, we generated reciprocal BM chimeras **(Figure 3G)**. The results confirmed that *Fam83h*-deficiency in stromal cells, rather than immune cells, was responsible for BM lymphopenia and impaired T cell development (**Figure 3G**, right panels).

**Figure 3.**
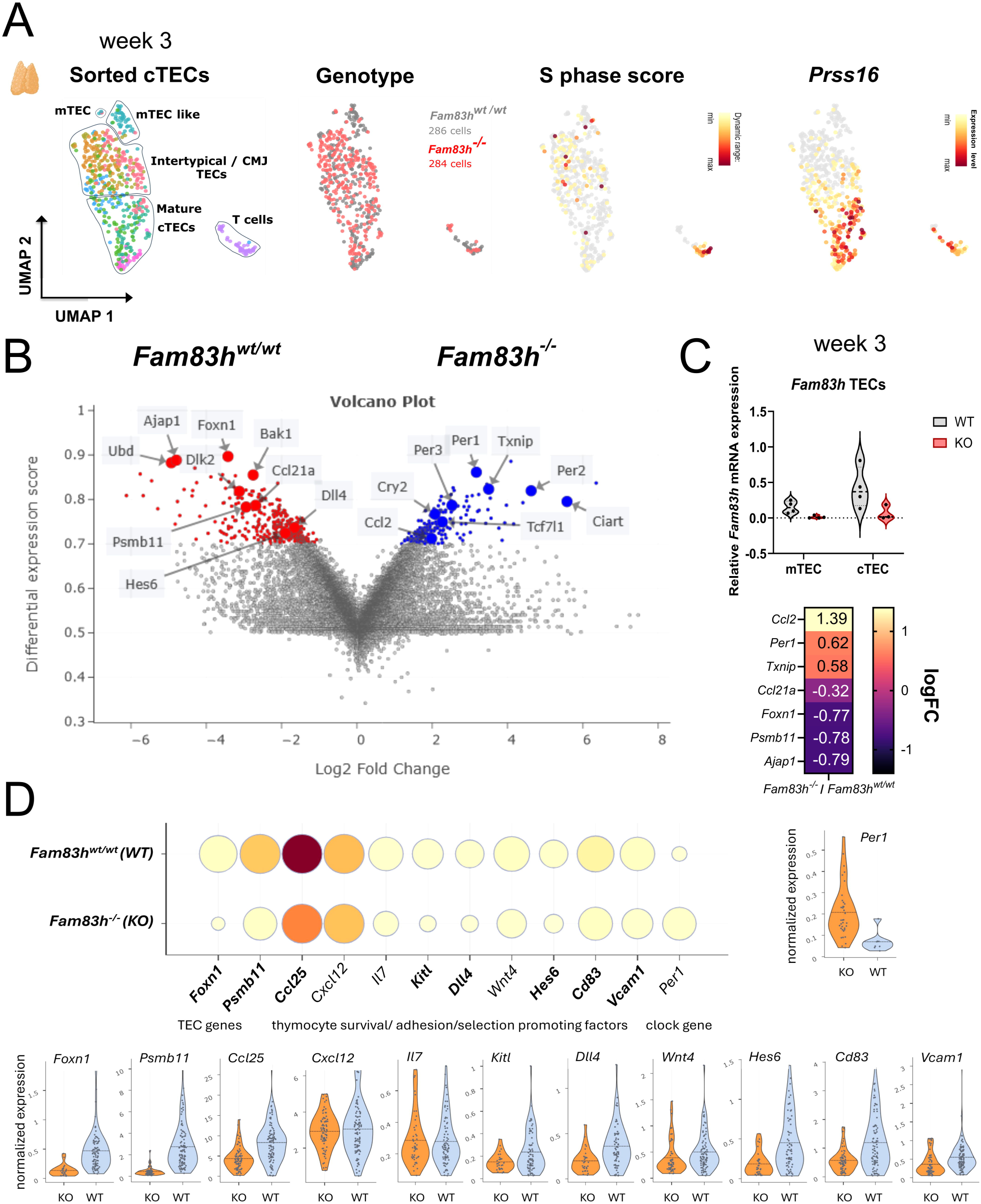
Thymocyte development is impaired in *Fam83h^-/-^* animals. (A) FCM analysis of the main thymocyte subsets: DN (green), DP (gray), CD4⁺ SP (blue), and CD8⁺ SP (magenta) at different time points (week 1,/ 2/ 3). DN1-4 subsets are shown in the bottom of the panel (DN1 (green), DN2 (light blue), DN3 (dark blue), DN4 (purple). The gating strategy is shown in Figure S12. Bars represent mean ± SD. Data are based on 3 independent experiments, *n*=9 for *Fam83h^wt/wt^* and *n*=7 for *Fam83h^-/-^* (week 1), *n*=11 for *Fam83h^wt/wt^* and *n*=6 for *Fam83h^-/-^* (week 2), and *n*=8 for *Fam83h^wt/wt^* and *n*=6 for *Fam83h^-/-^* (week 3). (B) Absolute quantification of thymic subsets by FCM at week 2. Total thymocyte counts, CD5 T cells/ thymus and fractions of the main thymocyte subsets: DN (green), DP (gray), CD4⁺ SP (blue), and CD8⁺ SP (magenta); DN1 (green), DN2 (light blue), DN3 (dark blue), DN4 (purple) were determined by FCM). The gating strategy is shown in Figure S12. Bars represent mean ± SD. Data are based on 3 independent experiments, *n*=6 for *Fam83h^wt/wt^* and *n*=5 for *Fam83h^-/-^*, unpaired *t-test*. p-values: *P ≤ 0.05, ****P ≤ 0.0001. (C) Absolute quantification of thymic subsets by FCM at week 2. Total cell counts/ thymus were determined for the main thymic subsets. The gating strategy is shown in Figure S10. Data are based on 3 independent experiments, *n*=6 for *Fam83h^wt/wt^* and *n*=5 for *Fam83h^-/-^*, unpaired *t-test* (box and whiskers, median/ min to max). p-values: *P ≤ 0.05, **P ≤ 0.01. (D) Quantification of pre-selection and post-selection DP thymocytes by FCM at weeks 2 and 3. The gating strategy is shown in Figure S12. Bars represent mean ± SD. Data are based on 3 independent experiments, *n*=6 for *Fam83h^wt/wt^* and *n*=5 for *Fam83h^-/-^* (week 2), Unpaired *t-test*. p-values: ns (P > 0.05) (E) Proliferation was determined by FCM analysis of Ki67 levels in thymocyte subsets (Figure S13). Representative histograms showing the decrease of Ki67 expression in *Fam83h^-/-^*(red) as compared to *Fam83h^wt/wt^* (gray) DN3 thymocytes at week 2-3 of age. The isotype control signal is depicted by the dashed line. The plot below shows the quantification of Ki67 staining in DN1–DP3 T cell subsets. Data are presented as min-max normalized percentages, *n*=4 for *Fam83h^wt/wt^* and *n*=4 for *Fam83h^-/-^*, unpaired *t-test*, p-values: ns (P > 0.05), *P ≤ 0.05, **P ≤ 0.01, ***P ≤ 0.001. (F) Percentage of late apoptotic cells among developing T cell subsets were determined by FCM at week 3. The gating strategy is shown in Figure S12. Data are based on 2 independent experiments, *n*=5 for *Fam83h^wt/wt^* and *n*=3 for *Fam83h^-/-^*, unpaired *t-test* (box and whiskers, median/ min to max). p-values: ns (P > 0.05).(G) Schematic diagram illustrating the generation of *Fam83h^-/-^* and *Fam83h^wt/wt^* BM chimeras. FCM analysis of thymus (upper middle panel, Figure S14) and blood (upper right panel, Figure S15) and bone marrow (lower middle and right panels, Figure S16) was performed at week 3–4 post-BM transfer to assess the contribution of donor-derived hematopoietic stem cells (HSCs) to the main hematopoietic lineages. Quantification of DN (green), DP (gray), CD4⁺ SP (blue), and CD8⁺ SP (magenta) thymocytes (Middle panel) and hematopoietic lineages in the peripheral blood. Data are based on 5 independent experiments, *n*=7 from *Fam83h^wt/wt^* to *Fam83h^wt/wt^, n*=7 from *Fam83h^wt/wt^* to *Fam83h^-/-^, n*=12 from *Fam83h^-/-^*to *Fam83h^wt/wt^* (blood)*, n*=12 from *Fam83h^wt/wt^* to *Fam83h^wt/wt^, n*=4 from *Fam83h^wt/wt^* to *Fam83h^-/-^, n*=16 from *Fam83h^-/-^* to *Fam83h^wt/wt^* (thymus). Bars indicate mean ± SD. Development of B and NK cells in the BM of reciprocal BM chimeras. Frequencies of B and NK cell progenitors in donor-derived BM were determined at week 3–4 post-transplantation. Bars indicate mean ± SD. Sample sizes: for B cells, *n*=4, 7, 9 for WT→WT, WT→KO, and KO→WT groups, respectively; for NK cells, *n*=3, 7, 6 for WT→WT, WT→KO, and KO→WT groups, respectively (NK; natural killer, R-NKP; refined natural killer progenitors).

Together, the reduced thymic output observed in *Fam83h^-/-^* animals is therefore more consistent with insufficient proliferation in the context of *Fam83h*-deficient stroma, rather than a block in selection processes or increased apoptosis rate.

### 2.6 Fetal hematopoiesis is intact in *Fam83h^-/-^* animals

As numerous thymic deficiencies arise during embryogenesis, we analyzed hematopoiesis in fetal liver (FL) and thymus at embryonic days E17.5 and E18.5 (29). No differences were observed in FL hematopoiesis and early T cell development was also intact at E17.5 (**Figure S5A**). Imaging techniques, including µCT and IF imaging at E18.5 and E17.5 respectively, revealed normal thymus formation, positioning, size and thymic architecture in *Fam83h^-/-^* embryos compared to *Fam83h^wt/wt^* littermates (**Figures S5B, C**).

### 2.7 Single-cell profiling reveals FAM83H-dependent regulation of cTEC identity

To assess the molecular impact of *Fam83h* deficiency in TECs, we performed single-cell RNA sequencing (scRNA-seq) on sorted cortical TECs (cTECs) from *Fam83h^-/-^* and *Fam83h^wt/wt^* mice at postnatal week 3. The sc-transcriptomic analysis confirmed the previously described heterogeneity of week 3 old cTECs, sorted as CD45^−^EpCAM^+^UAE^−^Ly.51^+^ cells (20, 21). The identified clusters included mature cTECs, (expressing *Prss16, Psmb11, Krt8, Foxn1, Ccl25*), proliferative intertypical/ cortico-medullary junction TECs, which shared some markers of perinatal (*Syngr1, Cxcl12*) and intertypical TECs (*Pdpn, Ly6a*), and showed higher S-phase score, and two smaller clusters expressing makers of mTECs. We assigned these two clusters as an mTEC-like cluster enriched in *Ccl21a* and *Krt5* and a minor subset of a few cells expressing *Aire*, likely representing mature mTECs. Additionally, a minute cluster of contaminating T cells was identified based on hematopoietic marker gene expression (*Cd3e*) **(Figures 4A, and S6A).**

**Figure 4.**
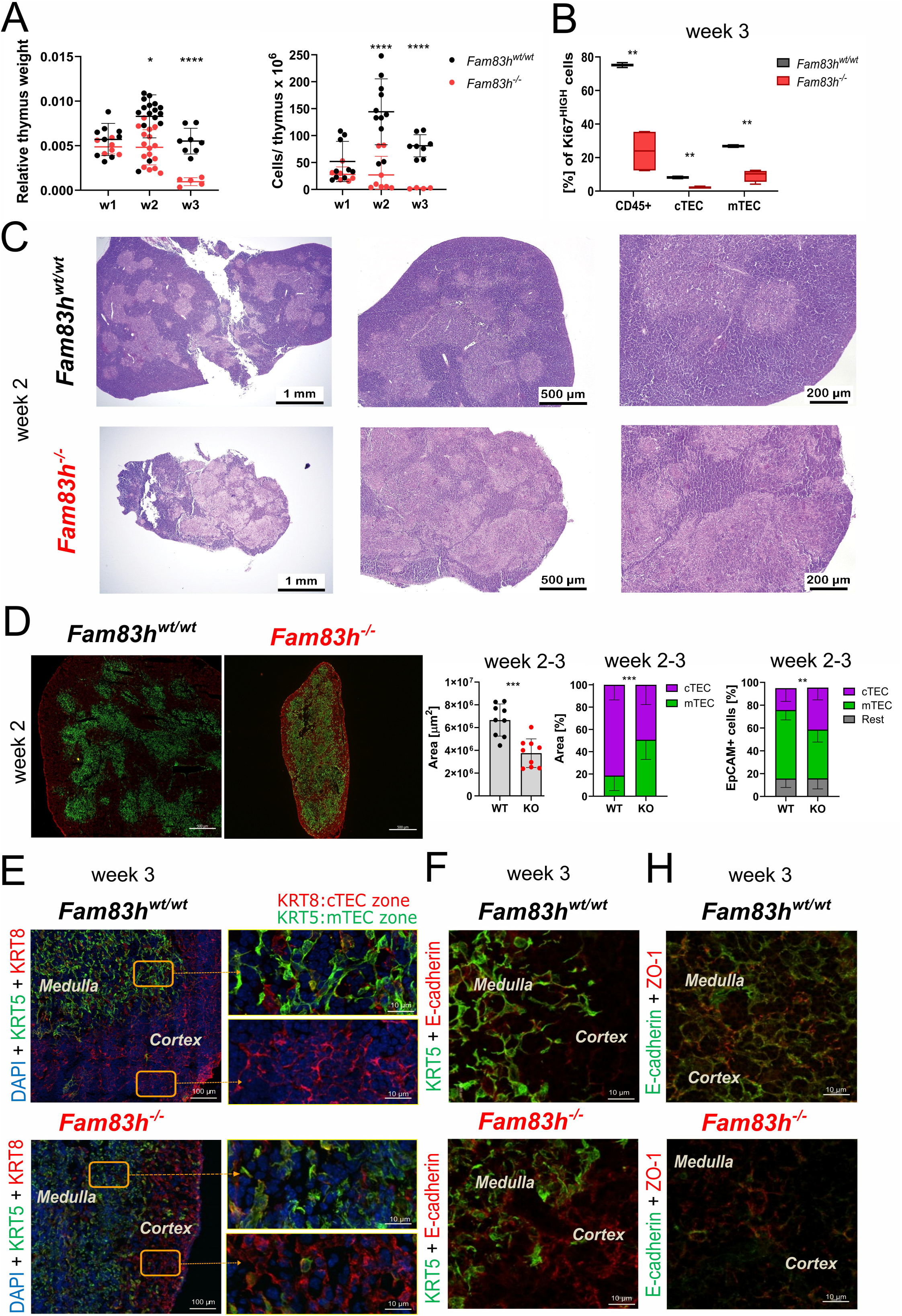
*Fam83h* deficient cTECs fail to express *Foxn1* and its target genes. (A) EpCAM+UAE-Ly51+ cTECs were single-sorted (Figure S17) from *Fam83h^wt/wt^* and *Fam83h^-/-^* thymi at 2 weeks of age and processed by sc-RNAseq using Smart-seq technology (42) UMAP projection shows 570 cells that passed quality control. Leiden clusters were identified using the Scarfweb platform and manually annotated (far left panel). Colors indicate unsupervised graph-based transcriptomic clusters, while black contour lines represent manually assigned TEC subset annotations. UMAP projections show an overlay of *Fam83h^wt/wt^* (gray, 286 cells) and *Fam83h^-/-^*(red, 284 cells) cells (left panel), S phase score (right panel), and cTEC marker gene, *Prss16*, expression level (far right panel). (B) Genes differentially expressed by mature cTECs isolated from *Fam83h^wt/wt^* and *Fam83h^-/-^* thymi. (C) RT-qPCR analysis of *Fam83h* mRNA expression in *Fam83h^wt/wt^* and *Fam83h^-/-^*mTECs and cTECs (upper panel, violin plot (median) n=3-4) and of selected genes in *Fam83h^wt/wt^* and *Fam83h^-/-^* cTECs (bottom panel). *Casc3* was used as reference gene. The heatmap displays log fold change in *Fam83h^-/-^* / in *Fam83h^wt/wt^*. (D) Expression of selected TEC identity, thymocyte survival, adhesion, and selection promoting factors and circadian rhythm gene *Per1* in *Fam83h^wt/wt^* and *Fam83h^-/-^* cTECs.

We focused our differential expression analysis on the mature cTEC cluster enriched in *Foxn1* as well as its multiple target genes **(Figures 4B and S5B)**, *Fam83h^wt/wt^* cTECs (red) expressed key TEC regulators, including *Foxn1, Ccl21a, Psmb11, Ubd, Dll4, Bak1,* as well as genes involved in cell adhesion (*Ajap1*) and Wnt or Notch signaling (*Tcf7, Dlk2, Hes6*). In contrast, *Fam83h^-/-^* cTECs (blue) were enriched for transcripts involved in circadian rhythm regulation, such as *Per1, Per2, Per3, Cry1, Cry2, Clock*, and *Ciart* **(Figure S6C)**. Among 338 genes enriched in *Fam83h^wt/wt^* cTECs (Nygen score >0.7, log2FC >1), 29 were identified as high-confidence *Foxn1* targets (from 450 genes in total, (30)) suggesting that *Fam83h*-deficient cTECs cannot activate the *Foxn1* transcriptional program efficiently and ultimately fail to differentiate. RT-qPCR analysis of sorted cTECs (*Fam83h^-/-^*/*Fam83h^wt/wt^*) confirmed the downregulation of *Foxn1* (logFC; -0.77), *Psmb11* (-0.78), *Ajap1* (-0.79), and *Ccl21a* (-0.32) in *Fam83h^-/-^* cells **(Figure 4C)**. Conversely, *Ccl2, Per1,* and *Txnip* were upregulated, potentially reflecting alterations in TEC homeostasis, chemotaxis, circadian rhythms and oxidative stress, validating the scRNA-seq findings on bulk cTEC samples **(Figure 4C)**. We have also analyzed expression levels of thymocyte survival or selection promoting factors and adhesion and signaling molecules in cTECs from *Fam83h^wt/wt^* and *Fam83h*-deficient mice. Among them, *Ccl25, Kitl, Dll4, Wnt4, Cd83, Vcam1* and *Hes6* were decreased in the absence of FAM83H (**Figure 4D**).

Overall, these results demonstrate that FAM83H is critical for TEC development and function, as its absence leads to disruption of the *Foxn1*-dependent cTEC transcriptional program.

### 2.8 FAM83H is essential for CK1 association with the keratin cytoskeleton

Given the established role of CK1α (*Csnk1a1*) in keratin cytoskeleton organization (1) we examined whether this pathway might also be affected in our model. We observed disrupted keratin-5 and keratin-8 architecture in *Fam83h^-/-^*TECs, along with broader epithelial defects in both, *Fam83h^-/-^* and *Fam83h*^Δ*87/*Δ*87*^ animals. These findings prompted us to investigate whether the truncated FAM83H protein, which lacks the first 87 amino acids in its N terminal domain, retains the ability to bind toCK1α. Due to the scarcity of TECs, we used ileum epithelial lysates from *Fam83h^wtwt,^ Fam83h^-/-,^* and *Fam83h*^Δ*87/*Δ*87*^ animals (2-3 weeks old) for co-immunoprecipitation (Co-IP) with anti-CK1 and anti-FAM83H antibodies, followed by mass spectrometry (MS) analysis. As shown in Figure 1B, ileum shows abundant expression of *Fam83h*. Co-IP with anti-FAM83H antibody confirmed the FAM83H–CK1α interaction in *Fam83h^wt/wt^* samples, which was absent in *Fam83h^-/-^* lysates **(Figure 5A).**

**Figure 5.**
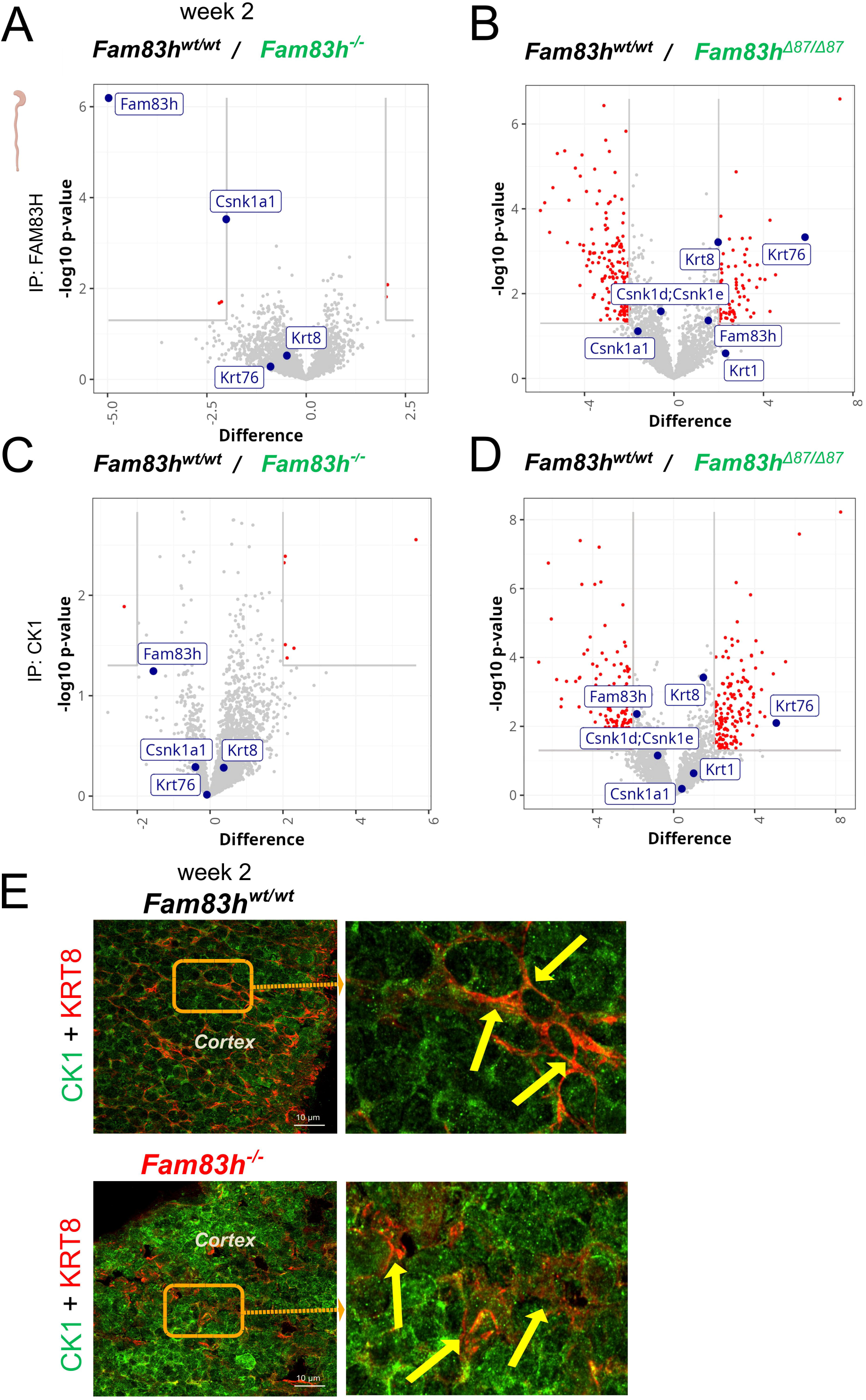
Mass spectrometry analysis of FAM83H and CK1 interacting proteins in ileum tissue from *Fam83h^wt/wt^, Fam83h^-/-,^* and *Fam83h*^Δ*87/*Δ*87*^ *confirms that Fam83h*^Δ*87/*Δ*87*^ fails to interact with CK1a. (A-D) Mass spectrometry was performed on ileum lysates immunoprecipitated from *Fam83h^wt/wt^, Fam83h^-/-^*, and *Fam83h*^Δ*87/*Δ*87*^ samples using anti-CK1 and anti-FAM83H antibodies. The analysis aimed to identify FAM83H and CK1-associated proteins and compare their abundance between *Fam83h^wt/wt^* and Fam83h deficient (*Fam83h^-/-^*) (A, C) and *Fam83h truncated (Fam83h*^Δ*87/*Δ*87*^*) samples (B, D).* Data represents 5 *Fam83h^wt/wt^,* 4 *Fam83h^-/-^* and 6 *Fam83h*^Δ*87/*Δ*87*^. (E) Cryosections from week 2 old *Fam83h^wt/wt^ and Fam83h^-/-^* thymi were stained for CK1α (green) and keratin-8 (red) were imaged by SDCM. Krt8+ CK1α+ TECs are marked by yellow arrows. Representative images from 3 independent experiments. Scale bar = 10 μm.

Additionally, samples from *Fam83h*^Δ*87/*Δ*87*^ mice showed a substantially reduced FAM83H–CK1 interaction **(Figure 5B).** Co-IP using CK1α antibody corroborated these findings: the FAM83H–CK1α interaction was seen in *Fam83h^wt/wt^* lysates but not in those from *Fam83h^-/-^* **(Figure 5C)** and *Fam83h*^Δ*87/*Δ*87*^ samples **(Figure 5D).**

IF imaging of thymic cortex revealed that in *Fam83h^wt/wt^*animals, CK1α localized to well defined vesicular structures, evenly distributed in the proximity of the cell membrane. In contrast, in the absence of *Fam83h*, CK1α distribution was more diffuse or aggregated in larger structures **(Figure 5E)**. Taken together, loss or mutation of FAM83H disrupts the CK1-keratin cytoskeletal network, potentially contributing to epithelial structural abnormalities and signaling defects.

## 3. Discussion

To date, FAM83H has primarily been studied in the context of amelogenesis imperfecta AI (3) intracellular transport, cytoskeletal organization (1), and cancer (12). In this study, we uncovered a previously unrecognized role of FAM83H in lymphopoiesis and T cell development. Our data identify *Fam83h* as a critical regulator of epithelial integrity, thymic stroma architecture, TEC maturation, and T cell development, acting through its scaffolding function with CK1 and keratin cytoskeletal proteins. We confirmed *Fam83h* expression across multiple epithelial tissues, with unexpectedly high expression in the intestine, surpassing that in ameloblasts, where it was originally characterized (4). Surprisingly, *Fam83h* expression was also detected in hematopoietic organs such as the spleen and BM, suggesting potential roles beyond classical epithelial biology.

The early lethality and widespread epithelial defects observed in *Fam83h*-deficient mice underscore its essential role in maintaining epithelial homeostasis, consistent with earlier findings (27, 28). These findings align with the report from Zheng *et al.*, who described perinatal lethality, skin lesions, inflamed digits, and reduced size in *Fam83h* mutant mice (2). However, the precise cause of lethality in *Fam83h^-/-^* mice remains unclear.

Notably, we did not observe any dental abnormalities, consistent with findings by Wang *et al.* (31). However, altered bone mineralization suggests potential structural and developmental roles for FAM83H, supporting its known involvement in epithelial-mesenchymal interactions.

Unexpectedly, *Fam83h* deficiency also caused pronounced changes in the hematopoietic system, which had not been shown before. Peripheral blood analysis revealed a skewing toward the myeloid lineage and reduced lymphoid output, at 3 weeks of age. BM analysis demonstrated a pronounced reduction in common lymphoid progenitors and their B and NK cell progeny, implicating the role of *Fam83h* in early lymphoid lineage commitment. One of the most striking findings was the severe thymic atrophy in *Fam83h*-deficient mice. *Fam83h*-deficient mice exhibited reduced thymus size, disrupted stromal architecture, and impaired T cell output. TECs displayed altered morphology, reduced proliferation and a failure to support proper thymocyte development. FCM analysis revealed a significant reduction in DP thymocytes largely driven by defective proliferation of DN progenitors, particularly at the DN3–DN4 stages, confined to the cortical subcapsular zone. This suggests that loss of FAM83H in cTECs is the primary driver of the thymic phenotype. Reciprocal BM chimera experiments confirmed the stromal origin of the hematopoietic defects, indicating that FAM83H functions in a non-hematopoietic, niche-dependent manner to support immune development.

Interestingly, fetal hematopoiesis, thymic organogenesis and stromal architecture appeared intact in *Fam83h-*deficient embryos, indicating that *Fam83h* is dispensable during embryogenesis but becomes essential postnatally, when the T cell production is boosted in the mouse. We hypothesize that in the postnatal thymus when the demand for robust T cell output increases following massive BM progenitor seeding, FAM83H becomes essential for sustaining epithelial function (32).

Single-cell RNA sequencing of TECs further revealed a loss of mature cTEC identity in *Fam83h*-deficient mice. Canonical cTEC genes critical for early thymocyte survival, adhesion, and development—such as *Ccl25, Kitl, Dll4,* and *Cd83*—were downregulated, whereas transcripts associated with circadian rhythm and stress responses were upregulated (33–35). In the absence of FAM83H, cTECs failed to express *Ccl25* and *Kitl (Scf),* essential for recruitment and survival of early thymic progenitors, as well as *Dll4*, which drives T cell development. In contrast, *Cxcl12* and *Il7* expression was preserved, allowing thymocyte migration, differentiation, and survival to proceed to some extent. Notably, *Ccl25, Dll4*, and *Cd83* are directly regulated by *Foxn1*, which was among the most downregulated genes in *Fam83h*□*/*□ cTECs (30, 33).

Importantly, *Fam83h*□*/*□ cTECs exhibited increased mRNA levels of multiple core circadian clock genes, whose expression can be indirectly regulated by casein kinases 1δ and 1ε (36, 37). Since FAM83H binds not only CK1α but also CK1δ and CK1ε, the altered circadian gene expression may result from dysregulated casein kinase activity in the absence of FAM83H (5).

Additionally, expression of genes involved in Wnt and Notch signaling pathways was impaired (*Wnt4, Dll4, Hes6*), suggesting that FAM83H may influence TEC maturation through these key developmental pathways. Prior studies have demonstrated that FAM83H interacts with CK1 and modulates Notch signaling, further supporting this hypothesis, and linking these pathways (38, 39).

Collectively, our data suggest that FAM83H is not only essential for epithelial cytoskeletal organization (8) but also for sustaining the transcriptional program necessary for TEC identity and function (30). Although *Fam83h* has not been classified as a core thymic gene, its expression pattern across epithelial datasets such as those by Bornstein et al. indicates a significant role in TEC biology (20). Complementing this, Park *et al*. demonstrated that shifts in TEC populations correlate with stages of T cell development, initially dominated by cTECs and later balancing between cTECs and mTECs as thymocyte maturation proceeds. This observation supports the model of thymic cross-talk, wherein reciprocal signaling between developing thymocytes and TECs promotes their mutual differentiation and function (19). Given that *Foxn1* expression is regulated by Wnt signaling (40), it is plausible that FAM83H indirectly modulates key pathways governing TEC function through Wnt pathway regulation.

We confirm that FAM83H is essential for the association between CK1α and keratin cytoskeletal proteins, consistent with previous findings (38). The loss of this interaction in *Fam83h***-**deficient and Δ87 mutant mice likely contributes to structural defects in epithelial cells, including TECs.

These data also suggest that CK1α activity may be essential for hematopoiesis in the context of *Fam83h* deficiency. Previous studies have shown that altering Wnt/β-catenin signaling in thymic epithelium impairs embryonic thymus development (41), and it has been proposed that *Fam83h* deficiency may perturb Wnt/β-catenin signaling in epithelial cells (24). Future studies should investigate whether *FAM83H* mutations contribute to epithelial or immune pathologies in human diseases, particularly those with congenital or early-onset manifestations. In conclusion, our study establishes *Fam83h* as a key regulator of thymic epithelial cytoskeletal integrity and a critical determinant of thymic and hematopoietic homeostasis. Through its role in scaffolding CK1 and organizing the keratin cytoskeleton, FAM83H supports the structural and functional competence of epithelial tissues, particularly those governing immune development. Its loss leads to widespread defects in epithelial morphology, signaling, and lymphoid cell production, offering new insights into congenital disorders with both epithelial and immune components.

## 4. Data Limitations and Perspectives

Our BM chimera experiments confine the functional role of FAM83H to the stroma (not necessarily restricted to the thymus) and exclude a role for FAM83H within hematopoietic cells. To definitively establish a TEC-intrinsic function of FAM83H, conditional TEC-specific deletion would be required. Nonetheless, multiple independent lines of evidence-including expression, structural, functional, and transcriptomic data-strongly point to a TEC-intrinsic requirement. Future studies employing either TEC-specific FAM83H knockout mice or thymic transplantation approaches could be essential to define the precise role of FAM83H in TECs.

Due to technical constraints-including the very low TEC yield per thymus, the risk of thymocyte contamination, the requirement for enzymatic digestion, and the limited availability of *Fam83h*^□*/*□^ thymi-we performed the co-IP experiment followed by mass spectrometry using ileal lysates to demonstrate that CK1 binds WT, but not Δ87FAM83H. The observation that this interaction is lost in the Δ87 variant-and that both *Fam83h*^□*/*□^ and *Fam83h*^Δ*87/*Δ*87*^ mice display nearly identical phenotypes-strongly supports the critical role of the CK1–FAM83H interaction. Since TECs clearly express both Fam83h and CK1α, we propose that the CK1–FAM83H interaction observed in ileal samples may also occur in cTECs.

## 5. Materials and Methods

### 5.1 Animals

C57BL/6 mice were housed in individually ventilated cages (Tecniplast) with free access to water and food (Altromin), Safe Select Fine bedding (Velaz), and environmental enrichment under a 12/12 light-dark cycle. The study complied with EU laws (Project licence AVCR 4212/2023 SOV II). Experiments followed the Czech Centre for Phenogenomics (CCP) Standard Operating Procedures.

#### Generation of *Fam83h*^Δ*87/*Δ*87*^ and *Fam83h^-/-^*mice

In this study, *Fam83h* mutant mice (*Fam83h*^Δ*87/*Δ*87*^) with a 119 bp deletion on exon 2 **(Figure S6A)** and *Fam83h* knockout (*Fam83h^-/-^*) mice (CRISPR/Cas9 technology) were generated within the IMPC effort (28). In *Fam83h^-/-^*, the whole protein-coding sequence was deleted **(Figure 1C)**. The location of gRNAs was guiding RNA 1 > 23 bp (7595–7617) and guide RNA 2 > 23 bp (12522–12544), which are located on exon 2 and exon 5, respectively, and were selected as the target site, are shown in **Figure 1C**. For genotyping, specific primers were designed (listed in **Supplementary Table 1**).

### 5.2 Relative mRNA Expression (RT-qPCR)

To investigate RNA levels in *Fam83h^wt/wt^* and *Fam83h^-/-^* mice for comparison, total RNA was extracted with Qiagen RNeasy Mini Kit (Cat# 74104) and Micro Kit (Cat# 74004), approximately 0.3 - 1 μg of RNA was used for cDNA synthesis (Thermo Fisher, RevertAid First Strand cDNA Synthesis Kit, Cat# K1621). Primers are listed in **Supplementary Table 2.**

### 5.3 Histological staining, Hematoxylin eosin (H&E), and immunofluorescence (IF)

#### 5.3.1 Hematoxylin eosin (H&E) staining

Paraffin sections were prepared by fixing samples in 4% PFA for >24 hours, transferring to 70% ethanol, embedding in cassettes, and sectioning at 4–5cμm (thymus) or 6–8cμm (paws). Forepaw samples were decalcified in Osteosoft (∼1 month). Slides were deparaffinized (Leica ST5010–ST5030), and antigen retrieval was performed in 10mM citric acid (pH 6) ∼15 min. H&E staining was done using Leica ST5020 + ST5030. Images were acquired on a Zeiss Axioscan.Z1 and Leica DM3000 LED microscopes at 20x/40x magnifications and analyzed using ZEN 3.1 and Leica Application Suite V4.13 software.

#### 5.3.2 Immunofluorescence (IF) staining

For cryosections, thymi were embedded in OCT, snap-frozen, sectioned at 7–10 μm at -20°C, and stored at -20°C. Slides were warmed ∼10-15 min at RT, and fixed in 4% PFA, permeabilized (0.2% Triton X, 10 min), blocked (2.5% BSA), and incubated with primary antibodies (listed in **Supplementary Table 3**) overnight. Secondary antibodies and DAPI staining were applied the next day. Slides were mounted with fluorescent mounting medium, dried, and imaged using Zeiss Axio Imager Z.2 and Nikon CSU-W1 confocal microscope equipped with a Prior automated stage controlled by NIS Element Software at 20x and 100x objectives. Confocal Z stacks were generated through the entire z-axis of thymus tissues. Image analysis was performed using NIS-Elements (with AI) 6.0.2.

#### 5.3.3 Quantitative analysis of cortical and medullary areas

Image segmentation and mask generation were performed in ImageJ (FIJI) v1.54p. Multichannel images (DAPI, red=cTEC, green= mTEC) were split and pre-processed (contrast equalization (0.1%), rolling-ball background subtraction (50 px), and Gaussian smoothing (σ=1)); mTEC received the same preprocessing except contrast enhancement. DAPI and cTEC were segmented using the Li algorithm, and mTEC was thresholded using Otsu method (for data from Nikon CSU-W1 Triangle method was used for thresholding). The total tissue ROI was defined by merging cTEC and mTEC masks, followed by hole filling, 3× dilation, and 3× erosion to close gaps. After 4-connected component labeling, only the largest object was retained and eroded by 5 px to remove edge artifacts. Logical AND operations restricted all masks (cTEC, mTEC, and their overlaps) to the ROI, and areas were measured for each.

### 5.4 Micro-computed tomography (µCT) analysis

Crania and paws were dissected and preserved in 4% PFA for at least 3 days, and scanned in Skyscan 1176 (Bruker, Belgium). NRecon 2.2.0.6 (Bruker, Belgium) was applied for reconstruction of virtual section. After fixation in 4% PFA, E18.5 embryos were stained with 25% Lugol’s solution for 10–14 days and embedded in 2.5% low-melting-point agarose (Sigma-Aldrich) one day before scanning. Scan was provided in Skyscan 1272 (Bruker, Belgium). InstaRecon 2.0.4.0 (InstaRecon, USA) plug-in was applied for reconstruction of the virtual section. Reconstructed data were visualized in CTvox (Bruker) with custom pseudocolor scales, and volumetric analysis was conducted using CT analyzer 1.18.4.0 (Bruker).

### 5.5 Flow cytometry (FCM) & Sorting

#### 5.5.1 Single cell suspension preparation

Thymocytes were released from thymi using mechanical dissociation through cell strainer, bone marrow (BM) cells were prepared by crushing femurs and tibias in mortar. Upon red blood cell (RBC) lysis using ACK, cells were resuspended in FACS buffer (2% FBS, 2mM EDTA, 10mM HEPES in HBSS without Ca^2+^Mg^2+^) and counted. For BM progenitor assays, RBC lysis and Fc blocking were omitted.

#### 5.5.2 Thymic stromal cell isolation

The thymus was enzymatically dissociated in RPMI + 2% FBS, 1 mg/mL Collagenase D, 1 mg/mL Dispase, and 40 U/mL DNase I, cut into small pieces, and incubated at 37°C for 30 minutes while rotating, with pipetting every 5 minutes. Afterward, FACS buffer (5–10mL) was added, and suspension was filtered through a >100µm mesh into 15mL tubes, adjusted to 5mL with FACS buffer. Then, RBCs were lysed, and cells were counted. CD45+ cells were depleted using EasySep™ Human CD45 Depletion Kit II (StemCells Technologies, Cat# 18945). After separation, cells were stained with the antibody mixture for 30 minutes on ice, then centrifuged and resuspended in FACS buffer.

#### 5.5.3 Blood sample processing

Blood samples were collected from the facial vein into EDTA-coated tubes. Following 2-3 rounds of RBC lysis, samples were centrifuged (400*g*, 10 min, 4°C). Cells were stained with a pre-mixed antibody cocktail containing Fc block for 20 minutes at 4°C, washed, and resuspended in a FACS buffer supplemented with counting beads. Immediately prior to analysis, Hoechst 33258 dye was added, and samples were acquired at high flow rate.

#### 5.5.4 Antibody staining - extracellular

Samples were stained with antibody cocktails (**Supplementary Table 4**) containing Fc block for 20 minutes at 4°C in the dark. Then, cells were washed with FACS buffer and centrifuged at 400*g* for 5 minutes at 4°C. The pellet was resuspended in FACS buffer. For the BM progenitor assay, containing CD16/32 antibody, no Fc block was added. For the apoptosis assay, cells were stained and washed in Annexin binding buffer instead of FACS buffer.

#### 5.5.5 Antibody staining - intracellular - Ki67 staining

Upon surface marker and viability staining (*Fixable Viability Dye eFluor™*), cells were fixed and permeabilized using the eBioscience Foxp3/transcription factor staining buffer set (Invitrogen #00-5523-00) according to the manufacturer’s instructions, washed, and stained with anti-Ki67 or isotype control antibody for 20-60 minutes at RT in the dark. Then, cells were washed with a permeabilization buffer.

#### 5.5.6 FCM data acquisition, sorting, and analysis

For absolute blood cell count analysis, 5 μL of counting beads (CountBright^TM^ Absolute, Invitrogen, C36950) were added and acquired on Cytek® Aurora (5L 16UV-16V-14B-10YG-8R, Cytek Biosciences). Sorting was performed with BD FACSAriaTM Fusion (4L 6V-2B-5YG-3R, BD Biosciences) in a biosafety Class II Type A2 cabinet. Full details of panels and gating strategies are described in **Figures S8-S18**. Data were analyzed using FlowJo software v10.10 (BD Life Sciences).

### 5.6 BM chimeras

P2-P3 neonates (CD45.2) were irradiated with two X-ray doses (each 3.5Gy) split by a 4-hour window. 5 × 10^6^ ACK lyzed whole BM cells from a congenic donor (CD45.1) were injected into liver in 25 μL of HBSS. Animals were observed daily, and engraftment was analyzed from peripheral blood at 3 weeks of age. Thymi and BM were analyzed for hematopoietic outcome at 3-4 weeks post-transplant.

### 5.7 Single-Cell RNA Sequencing

#### 5.7.1 Single cTECs sorting

Thymic stromal cells were isolated from week 2-old *Fam83h^wt/wt^*and *Fam83h^-/-^* animals (see above). cTECs were identified as CD45- Ter119- EpCAM+ UEA- Ly.51+ cells and single-cell sorted into 384-well plates already containing 0.3 µL of lysis buffer (29) and 3 µL of silicone oil (100 cSt, Sigma-Aldrich), using the Formulatrix Mantis. As a negative control, a subset of wells received no sorted cells.

#### 5.7.2 Generation of smart-seq3xpress libraries (scRNA-seq)

Smart-seq3xpress libraries were prepared following published protocol (42) involving reverse transcription and 14 cycles of PCR pre-amplification after sample storage at −80°C. Indexed libraries were generated by transferring cDNA into plates with dried primers, followed by Tn5 tagmentation and PCR amplification for 16 cycles. Libraries were purified using SeraMag and SPRIselect beads, quality-checked on a Fragment Analyzer, quantified by Qubit, and sequenced on an Illumina NextSeq 2000 with 2 × 109 bp paired-end and index reads.

#### 5.7.3 Analysis of scRNA-seq Data Alignment of Raw Data

The raw read files were processed with zUMIs (version 2.7.9e) (43), applying published parameters (42) and settings documented in the supplemental file (Smartseq3xpress_SS3x_manuscript_all.yaml). Reads were aligned with STAR (version 2.7.3a) against mouse genome GRCm38, using gencode.vM8 annotation. Several down sampled count matrices were created, and for downstream analyses, a range of 10,000–20,000 reads per cell was used.

### Data Processing

Subsequent steps were carried out in Seurat (version 5.2.1) (44), focusing on 5′-end tagged reads. Reads mapping to spike-ins (ERCC and SIRV) were excluded, and cells were filtered based on quality metrics (e.g., gene count >1200, RNA count >2200, spike-in <12%, mitochondria <6%). SCTransform with glmGamPoi was used for normalization (45, 46). Dimensionality reduction and clustering were done using the first 30 principal components, UMAP, and a resolution of 0.8. For integration, 50-dimension CCA via the IntegrateLayers function was applied. Seurat objects were converted to h5ad format and uploaded to Scarfweb. Data is available at https://www.nygen.io.

### 5.8 Mass spectrometry (MS) analysis of ileum samples

#### 5.8.1 Co-immunoprecipitation assay (Co-IP)

Dynabeads™ Protein G (Invitrogen, Cat# 10004D) and a specific lysis buffer (20mM Tris-HCl, pH 7.4; 150mM NaCl; 1% Triton X-100 with protease inhibitors) were used to immune-precipitate proteins from tissue lysates. Tissues were lysed using mechanical disruption and centrifuged, followed by pre-clearing with Dynabeads. Lysates were incubated with dynabeads coated with antibodies against Fam83h and CK1α (**Supplementary Table 5**), and immunoprecipitated proteins were processed for Western blot or MS. LC-MS analyses were conducted at the OMICS Core Facility, Biocev, Charles University (https://www.biocev.eu/en).

#### 5.8.2 Protein Digestion

Protein-bound beads were treated with TEAB/SDC buffer, reduced with TCEP, and alkylated with chloroacetamide. Proteins were digested on-bead with trypsin overnight at 37°C. After centrifugation and acidification with TFA, SDC was removed using ethyl acetate extraction (29). The resulting peptides were desalted using C18 stage tips (30).

#### 5.8.3 LC-MS 2 Analysis

LC-MS analysis was performed using a Nano reversed-phase column (C18) with mobile phase buffers A (water, 0.1% formic acid) and B (acetonitrile, 0.1% formic acid). Peptides were loaded onto a trap column for 4 minutes and eluted using a gradient from 4% to 35% B in 60 minutes. Peptides were ionized by electrospray ionization and analyzed on a Thermo Orbitrap Ascend. Survey scans of peptide precursors were conducted at 120K resolution, and tandem MS was carried out with HCD fragmentation. Ion count targets and dynamic exclusion were set for optimal analysis.

#### 5.8.4 Analysis of MS data

Data were analyzed and quantified using MaxQuant software (version 1.6.10.43) (31) with a false discovery rate (FDR) set to 1% for both proteins and peptides, and a minimum peptide length of seven amino acids. MS/MS spectra were searched using the Andromeda engine against a Mus musculus database. Specific modifications included carbamidomethylation of cysteine, acetylation of N-terminal proteins, and oxidation of methionine. The "match between runs" feature was applied for data transfer and quantification, utilizing label-free algorithms. Data was processed using Perseus 1.6.15.0, with log-transformed data and missing values imputed using a normal distribution method. For each antibody and corresponding dataset, t-tests were used to perform comparisons, and the results were visualized using a volcano plot. Genes showing at least a twofold difference and a statistically significant p-value (p < 0.05) were highlighted. All analyses were conducted in R version 4.4.3.

### 5.9 Statistical Analyses

Statistical analyses were conducted using GraphPad Prism (version 9). Data are presented as mean ± standard deviation (SD) or standard error of mean (SEM), with biological replicates (mice or embryos). Group comparisons were performed using unpaired two-tailed **Student’s t-test** for normally distributed datasets with two groups, for comparisons involving multiple groups or timepoints, **one-way** or **two-way ANOVA** was used. For FCM analyses, mean frequencies of cell subsets were compared across genotypes and developmental stages using ANOVA or t-tests, depending on experimental design. Significance was set at *p* ≤ 0.05. The level of significance is indicated in figures as follows: p-values: ns (P > 0.05), *P ≤ 0.05, **P ≤ 0.01, ***P ≤ 0.001, ****P ≤ 0.0001. In Smart-seq scRNA-seq analyses, cluster-based comparisons (e.g., S-phase scoring, gene expression) were performed using Nygen software pipeline (ScarfWeb) with in-built statistical modules (https://scarfweb.nygen.io/). Sample sizes, independent experiments, and statistical tests are reported in **figure legends**.

## Supporting information

Ogan_et_al_Supplementary-Information

## Data availability statement

The scRNA-seq data that support the findings of this study are openly available in Zenodo (doi: 10.5281/zenodo.15436243). The MS proteomics data that support the findings of this study are openly available in ProteomeXchange Consortium via the PRIDE (47) partner repository with the dataset identifier PXD064193.

## Ethics approval statement for animal studies

The study complied with EU laws (Project licence AVCR 4212-2023 SOV II). Experiments followed the Czech Centre for Phenogenomics (CCP) Standard Operating Procedures.

## Author contributions

Betul Melike Ogan, Veronika Forstlová, Laura Dowling, Kristína Vičíková, Kamila Křížová, Sylvie Červenková, František Špoutil, Michaela Procházková, Jolana Turečková, Olha Fedosieieva, Juraj Lábaj and Jana Balounová performed the experiments. Petr Nickl prepared the *Fam83h*-defficient mouse strains. Vendula Novosadová analyzed the data. Betul Ogan and Jana Balounová wrote the manuscript. Jana Balounová, Jan Procházka and Radislav Sedláček designed and supervised the study and obtained funding. All authors approved the final manuscript.

## Acknowledgments

This work was supported by *LM2023036* and **CZ.02.01.01/00/23_015/0008189** (*Upgrade of the large research infrastructure CCP III*), co-funded by the European Union and the Ministry of Education, Youth and Sports of the Czech Republic (MEYS). We also acknowledge previous support from the projects **LM2018126** (*Czech Centre for Phenogenomics*, MEYS) and **CZ.02.1.01/0.0/0.0/18_046/0015861** (*CCP Infrastructure Upgrade II*, MEYS and ESIF).

The authors acknowledge Imaging Methods Core Facility at BIOCEV, supported by the MEYS CR (**LM2023050** Czech-BioImaging), BIOCEV GeneCore Facility (https://www.biocev.eu/en/services/genecore-facility.7), and OMICS Mass Spectrometry Core Facility (Faculty of Science, Charles University) at Biocev research center for their support & assistance in this work.

The authors used OpenAI’s ChatGPT for grammar and language correction. All content and conclusions in this paper are the responsibility of the authors.

## Abbreviations

FAM83H: Family of Sequence Similarity 83H
CK1: casein kinase 1
TEC: thymic epithelial cells
scRNA-seq: single-cell RNA-sequencing
FCM: flow cytometry, bone marrow, BM
DP: the double-positive (CD4^+^CD8^+^)
DN: double negative (CD4^−^CD8^−^)
MS: mass spectrometry

