## Supplementary material for "FAM83H regulates postnatal T cell development through thymic stroma organization": Ogan_et_al_Supplementary-Information

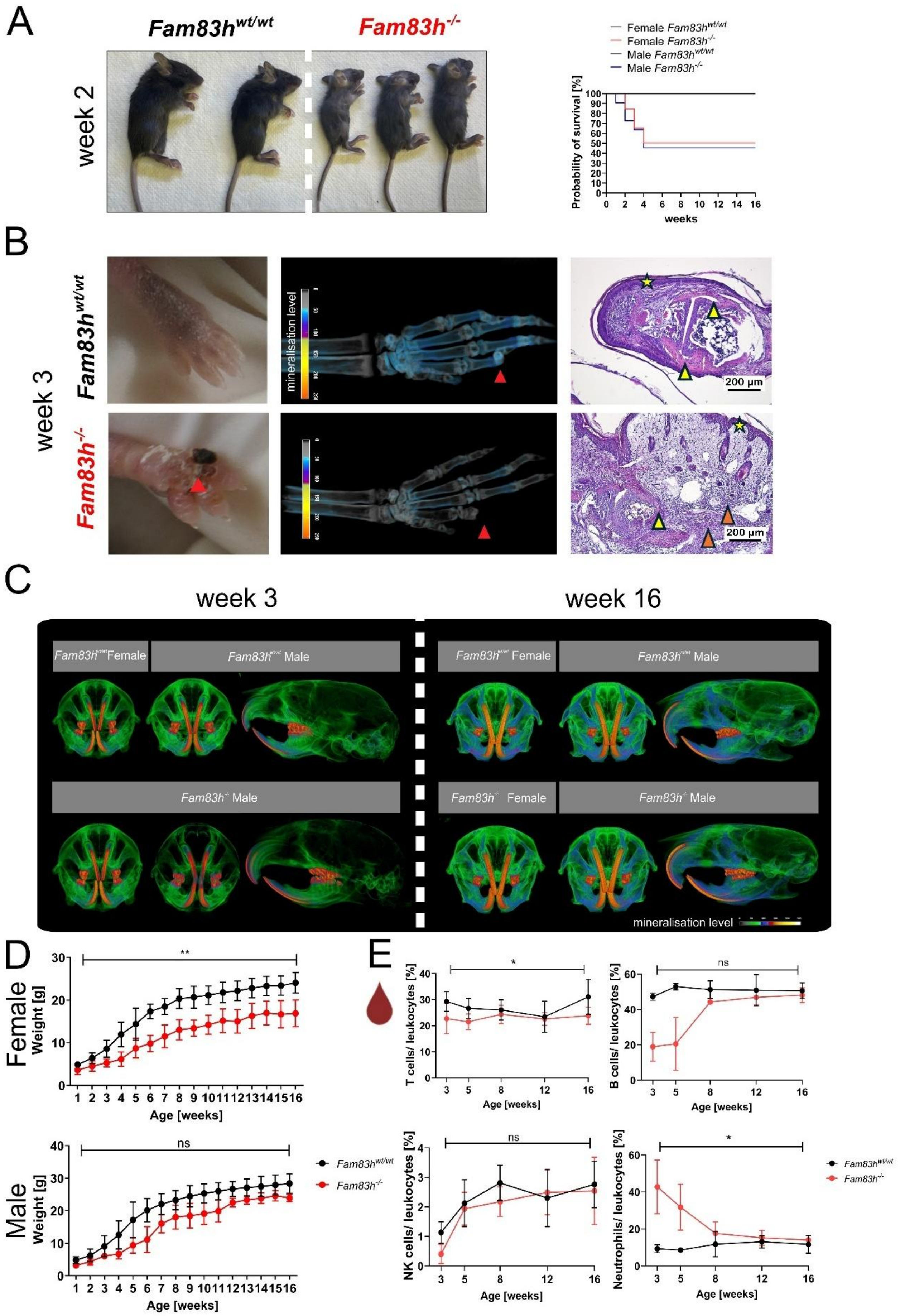
A
Fam83h wt/wt
Fam83h -/-
week 2
Female Fam83h wt/wt
Female Fam83h -/-
Male Fam83h wt/wt
Male Fam83h -/-
Probability of survival [%]
weeks
B
week 3
mineralisation level
200 µm
C
week 3
week 16
Fam83h wt/wt Female
Fam83h wt/wt Male
Fam83h -/- Male
Fam83h -/- Female
mineralisation level
D
Female
Weight [g]
Age [weeks]
Male
Fam83h wt/wt
Fam83h -/-
E
T cells/ leukocytes [%]
B cells/ leukocytes [%]
NK cells/ leukocytes [%]
Neutrophils/ leukocytes [%]
Age [weeks]
ns
*
**

**Supporting Information Figure 1 | Phenotypic characterization of *Fam83h*$^{wt/wt}$ and *Fam83h*$^{-/-}$ mice.**

(A) As compared to *Fam83h*$^{wt/wt}$, *Fam83h*$^{-/-}$ mice are smaller in size and show a disheveled coat at 2 weeks of age. The survival curve shown represents the probability of survival over 16 weeks in male and female mice with different *Fam83h* genotypes: *Fam83h*$^{wt/wt}$, *Fam83h*$^{-/-}$.

(B) At 3 weeks of age, *Fam83h*$^{-/-}$ mice show skin lesions (left panel), mineralization defects (middle panel, micro-computed tomography (µCT) imaging), and bone deformities (right panel, H&E staining). In H&E staining, vast infiltration of neutrophils, macrophages, and lymphocytes in *Fam83h*$^{-/-}$ mice (orange arrowheads), along with epithelial hyperplasia (yellow asterisk) and missing bone formation, connective tissue dysregulation, and cells are not differentiating correctly (yellow arrows) compared to *Fam83h*$^{wt/wt}$.

(C) Mineralization levels were analyzed in week 3 and week 16 of old animals using µCT imaging.

(D) Body weight progression – from week 3 to 16 of age. Points and connecting lines with error bars indicate mean ± SD. Statistical analysis was performed using a two-tailed unpaired *t-test*, p-values: ns ($P > 0.05$), $^{**}P \leq 0.01$. Sample sizes: 16 to 12 mice for male *Fam83h*$^{wt/wt}$ and *Fam83h*$^{-/-}$ groups, and 16 to 9 mice for female *Fam83h*$^{wt/wt}$ and *Fam83h*$^{-/-}$ groups, respectively.

(E) Leukocyte percentages in peripheral blood were measured using flow cytometry from week 3 to 16. Gating strategy is shown in **Figure S8**. Sample size n(*Fam83h*$^{wt/wt}$)=4-9, n(*Fam83h*$^{-/-}$)=7-12. Statistical analysis was performed using a two-tailed unpaired *t-test*, p-values: ns ($P > 0.05$), $^{*}P \leq 0.05$.

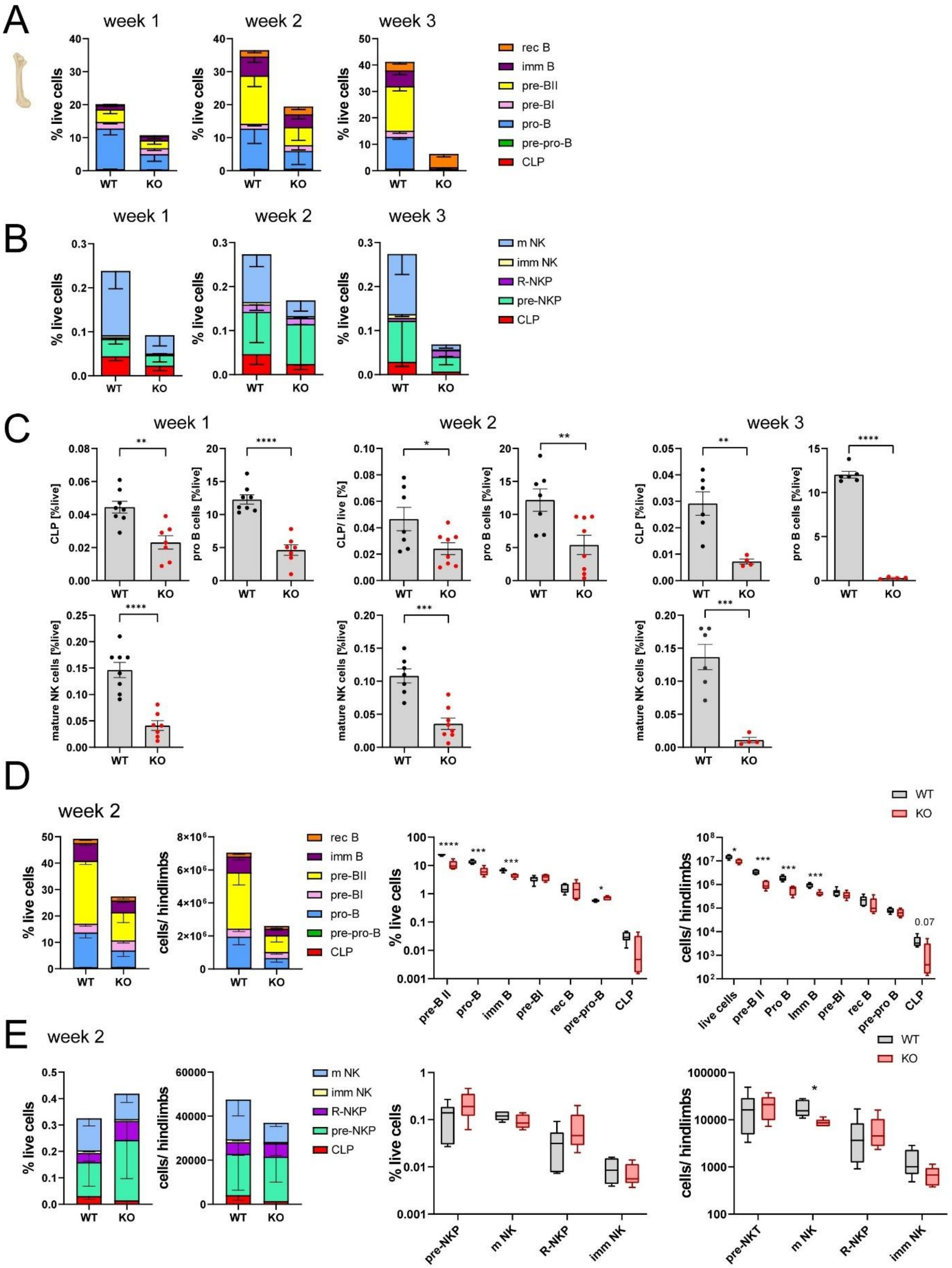
A
week 1
week 2
week 3
% live cells
WT
KO
rec B
imm B
pre-BII
pre-BI
pro-B
pre-pro-B
CLP
B
m NK
imm NK
R-NKP
pre-NKP
C
CLP [%live]
pro B cells [%live]
CLP/ live [%]
mature NK cells [%live]
D
week 2
cells/ hindlimbs
live cells
0.07
E
pre-NKT

**Supporting Information Figure 2 | Impaired B and NK cell development in the BM of *Fam83h$^{-/-}$* mice across early postnatal weeks.**

(A) B cell development in *Fam83h$^{wt/wt}$* and *Fam83h$^{-/-}$* mice from week 1 to week 3. Frequencies of B cell progenitors were determined in the BM of *Fam83h$^{wt/wt}$* and *Fam83h$^{-/-}$* mice by FCM. The full gating strategy is shown in **Figure S10**. Bars indicate mean ± SD. Sample sizes: *n*=8, 7 for *Fam83h$^{wt/wt}$* and *Fam83h$^{-/-}$* mice at week 1; *n*=7, 8 at week 2; *n*=6, 4 at week 3, respectively.

(B) NK cell development in *Fam83h$^{wt/wt}$* and *Fam83h$^{-/-}$* mice from week 1 to week 3. Frequencies of NK cell progenitors were determined in the BM of *Fam83h$^{wt/wt}$* and *Fam83h$^{-/-}$* mice by FCM. The full gating strategy is shown in **Figure S10**. Bars indicate mean ± SD. Sample sizes: *n*=8, 7 for *Fam83h$^{wt/wt}$* and *Fam83h$^{-/-}$* mice at week 1; *n*=7, 8 at week 2; *n*=6, 4 at week 3, respectively.

(C) Frequencies of common lymphoid progenitors (CLP), pro-B cells, and mature NK cells in *Fam83h$^{wt/wt}$* and *Fam83h$^{-/-}$* mice from week to week 3. The full gating strategy is shown in **Figure S10**. Symbols represent individual mice, and bars indicate mean ± SEM. Statistical analysis was performed using an unpaired *t-test,* p-values: *P ≤ 0.05, **P ≤ 0.01, ***P ≤ 0.001, ****P ≤ 0.0001. Sample sizes: *n*=8, 7 for *Fam83h$^{wt/wt}$* and *Fam83h$^{-/-}$* mice at week 1; *n*=7, 8 at week 2; *n*=6, 4 at week 3, respectively.

(D) Relative and absolute quantification of B cell progenitors at week 2. The full gating strategy is shown in **Figure S10**. Data are based on 3 independent experiments, n=6 for Fam83h$^{wt/wt}$ and n=5 for Fam83h$^{-/-}$ (box and whiskers, median/ min to max)*,* unpaired *t-test*. s: *P ≤ 0.05, ***P ≤ 0.001, ****P ≤ 0.0001.

(E) Relative and absolute quantification of NK cell progenitors at week 2. The full gating strategy is shown in **Figure S10**. Data are based on 3 independent experiments, n=6 for Fam83h$^{wt/wt}$ and n=5 for Fam83h$^{-/-}$ (box and whiskers, median/ min to max)*,* unpaired *t-test*. s: *P ≤ 0.05.

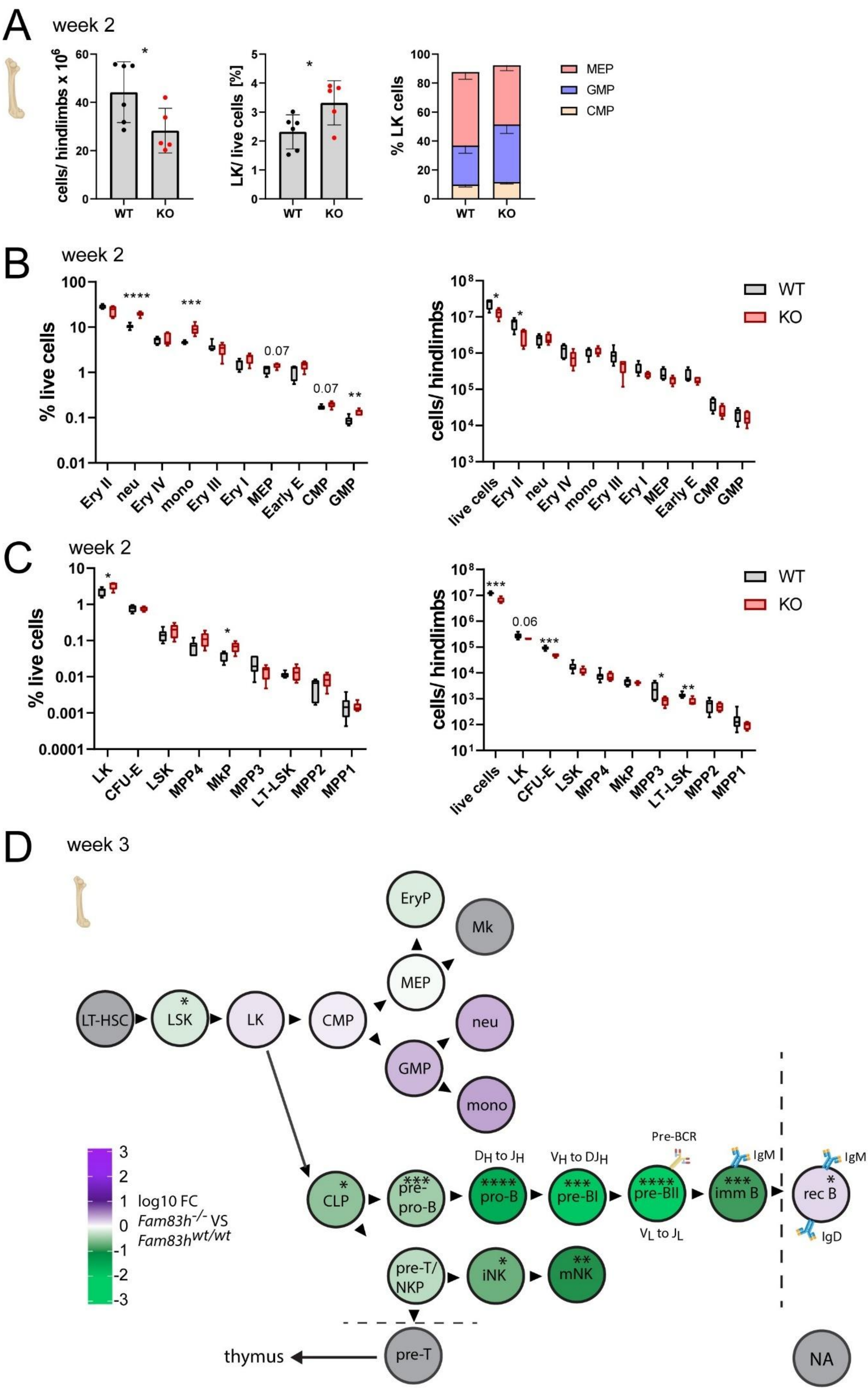
A
week 2
cells/ hindlimbs x 10^6
LK/ live cells [%]
% LK cells
WT
KO
MEP
GMP
CMP
B
week 2
% live cells
Ery II
neu
Ery IV
mono
Ery III
Ery I
MEP
Early E
CMP
GMP
0.07
cells/ hindlimbs
live cells
WT
KO
C
week 2
% live cells
LK
CFU-E
LSK
MPP4
MkP
MPP3
LT-LSK
MPP2
MPP1
0.06
cells/ hindlimbs
live cells
D
week 3
LT-HSC
LSK
LK
CMP
MEP
EryP
Mk
GMP
neu
mono
CLP
pre-pro-B
pro-B
pre-BI
pre-BII
imm B
rec B
D_H to J_H
V_H to DJ_H
V_L to J_L
Pre-BCR
IgM
IgD
pre-T/NKP
iNK
mNK
pre-T
thymus
NA
log10 FC
Fam83h-/- VS
Fam83hwt/wt

**Supporting Information Figure 3 | Erythro-myeloid and hematopoieitic stem cells and progenitors are not dramatically affected by *Fam83h*$^{-/-}$ deficiency.**

(A) Total BM cellularity and frequencies of Lineage$^{-}$ Kit$^{+}$(LK) progenitors and their progeny (common myeloid progenitors (CMP), granulocyte-macrophage progenitors (GMP), and megakaryocyte-erythrocyte progenitors (MEP)), were determined in the BM of *Fam83h*$^{wt/wt}$ and *Fam83h*$^{-/-}$ mice by FCM at week 2. The full gating strategy is shown in **Figure S9**. Bars indicate mean ± SD. Sample sizes: n=6, for *Fam83h*$^{wt/wt}$ and n=5 for *Fam83h*$^{-/-}$ mice, respectively. Statistical analysis was performed using an unpaired *t-test,* p-values: *P ≤ 0.05.

(B) Frequencies and absolute cell counts of erythro-myeloid progenitors and myeloid cells were determined in the BM of *Fam83h*$^{wt/wt}$ and *Fam83h*$^{-/-}$ mice by FCM at week 2. The full gating strategy is shown in **Figure S9**. Sample sizes: n=6, for *Fam83h*$^{wt/wt}$ and n=5 for *Fam83h*$^{-/-}$ mice, respectively (box and whiskers, median/ min to max)*,* unpaired *t-test*. s: *P ≤ 0.05, **P ≤ 0.01, ***P ≤ 0.001, ****P ≤ 0.0001.

(C) Frequencies and absolute cell counts of erythro-myeloid progenitors and myeloid cells were determined in the BM of *Fam83h*$^{wt/wt}$ and *Fam83h*$^{-/-}$ mice by FCM at week 2. The full gating strategy is shown in **Figure S11**. Sample sizes: n=6, for *Fam83h*$^{wt/wt}$ and n=5 for *Fam83h*$^{-/-}$ mice, respectively (box and whiskers, median/ min to max)*,* unpaired *t-test*. s: *P ≤ 0.05, **P ≤ 0.01, ***P ≤ 0.001.

(D) Schematic representation of BM progenitor development in BM in Fam83h-/- animals at week 3.

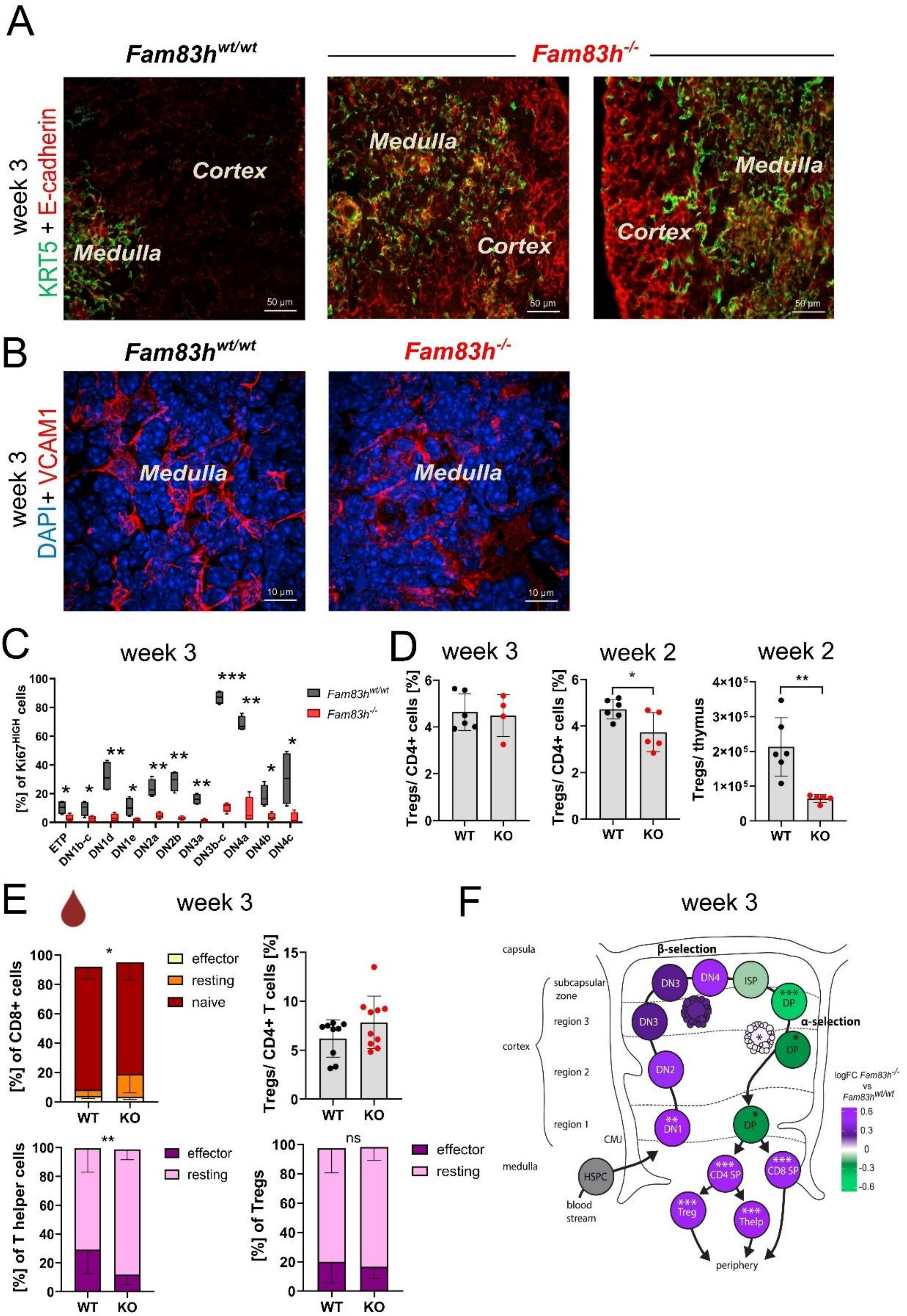
A
Fam83hwt/wt
Fam83h-/-
week 3
KRT5 + E-cadherin
Medulla
Cortex
50 µm
B
week 3
DAPI+ VCAM1
10 µm
C
week 3
[%] of Ki67HIGH cells
Fam83hwt/wt
Fam83h-/-
ETP
DN1b-c
DN1d
DN1e
DN2a
DN2b
DN3a
DN3b-c
DN4a
DN4b
DN4c
D
week 3
week 2
Tregs/ CD4+ cells [%]
Tregs/ thymus
WT
KO
E
week 3
[%] of CD8+ cells
effector
resting
naive
Tregs/ CD4+ T cells [%]
[%] of T helper cells
[%] of Tregs
ns
F
week 3
capsula
subcapsular zone
region 3
cortex
region 2
region 1
CMJ
medulla
β-selection
α-selection
DN3
DN4
ISP
DP
DN2
DN1
HSPC
blood stream
CD4 SP
CD8 SP
Treg
Thelp
periphery
logFC Fam83h-/- vs Fam83hwt/wt

**Supporting Information Figure 4 | *Fam83h$^{-/-}$* mice show altered thymic corticomedullary architecture and decreased T cell progenitor proliferation at week 3.**

(A) Cryosections from week 3 old *Fam83h$^{wt/wt}$ and Fam83h$^{-/-}$* thymi were stained for keratin-5 (green) and E-cadherin (red), and imaged with SDCM Nikon CSU-W1. Scale bar: 50 µm.

(B) Cryosections from week 3 old *Fam83h$^{wt/wt}$ and Fam83h$^{-/-}$* thymi were stained for VCAM1 (red) and nuclei were visualized by DAPI (blue), imaged with SDCM Nikon CSU-W1. Scale bar: 10 µm.

(C) Proliferation of DN thymocyte subsets. Frequency of Ki67HIGH cells in DN T cell subsets was analyzed by flow cytometry at week 3. Data are presented as min-max normalized percentages, *n*=4 for both *Fam83h$^{wt/wt}$ and Fam83h$^{-/-}$*, unpaired *t-test*, p-values: *P ≤ 0.05, **P ≤ 0.01, ***P ≤ 0.001.

(D) Percentage of Tregs out of CD4+ Tcells (week 3, week 2) and absolute Treg counts (week 2) were determined in thymi of *Fam83h$^{wt/wt}$ and Fam83h$^{-/-}$*animals.The full gating strategy is shown in **Figure S12**. Bars indicate mean ± SD. Sample sizes: *n*=6/6 for *Fam83h$^{wt/wt}$* and *n*=4/5 for *Fam83h$^{-/-}$* mice at weeks 3 and 2 respectively. Statistical analysis was performed using an unpaired *t-test,* p-values: *P ≤ 0.05, **P ≤ 0.01.

(E) Percentage of Tregs out of CD4+ Tcells and fractions of effector/ resting T cell subsets were determined in peripheral blood of *Fam83h$^{wt/wt}$ and Fam83h$^{-/-}$*animals at week 3. The full gating strategy is shown in **Figure S8.** Bars indicate mean ± SD. Sample sizes: *n*=9 for *Fam83h$^{wt/wt}$* and *n*=10 for *Fam83h$^{-/-}$* mice at weeks 3 and 2 respectively. Statistical analysis was performed using an unpaired *t-test,* p-values: *P ≤ 0.05, **P ≤ 0.01.

(F) Schematic representation (modified from (Blackburn and Manley 2004)) of developmental blocks in T cell development in *Fam83h$^{-/-}$* animals based on FCM analysis of week 3 old thymi (**Figure S12**). LogFC *Fam83h$^{-/-}$* vs *Fam83h$^{wt/wt}$* is shown for each T cell stage. Statistical analysis was performed using an unpaired *t-test,* p-values: *P ≤ 0.05, **P ≤ 0.01, ***P ≤ 0.001.

*Blackburn, C. C. and N. R. Manley (2004). "Developing a new paradigm for thymus organogenesis." Nature Reviews Immunology **4**(4): 278-289.*

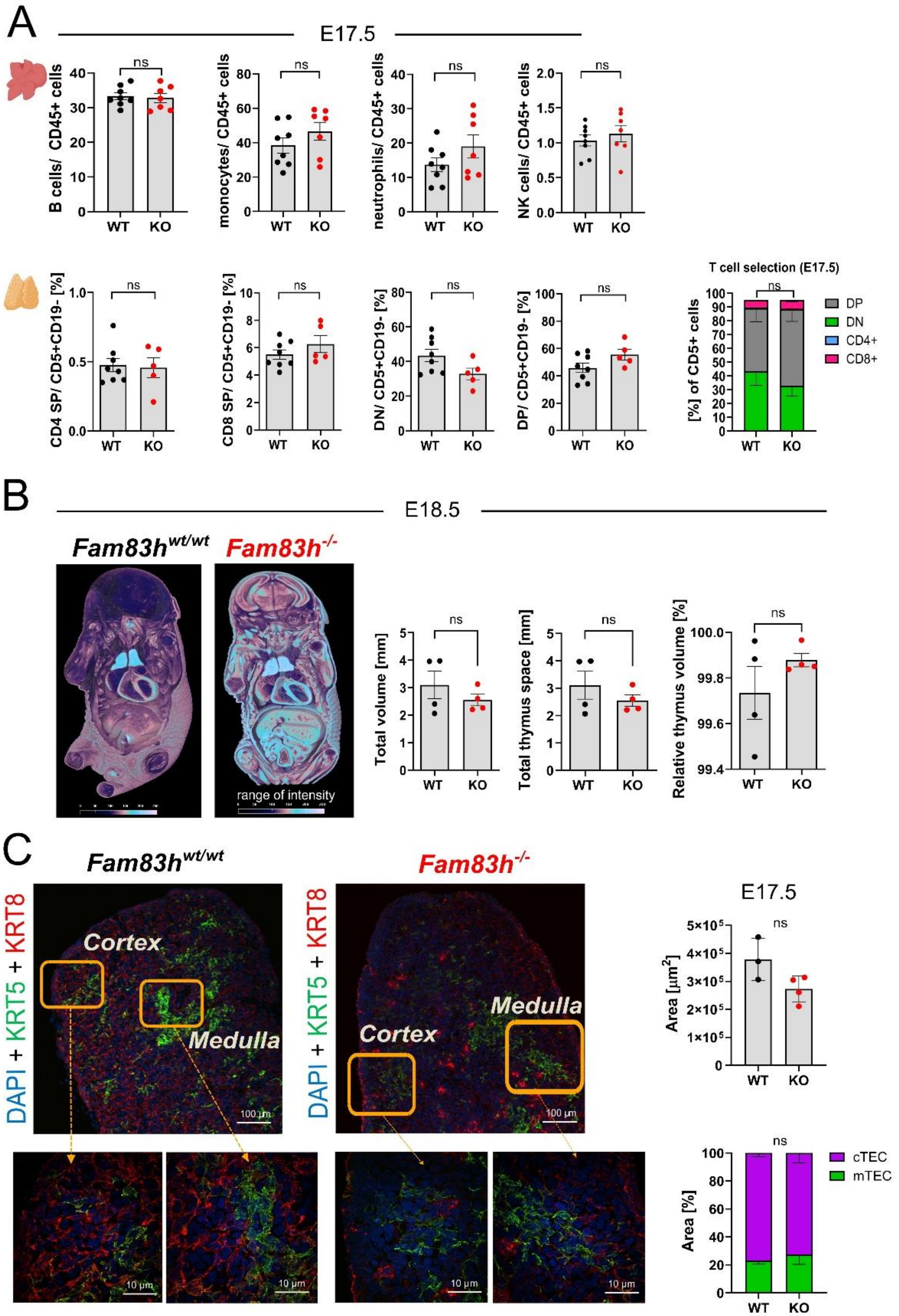

A
E17.5
B cells/ CD45+ cells
monocytes/ CD45+ cells
neutrophils/ CD45+ cells
NK cells/ CD45+ cells
CD4 SP/ CD5+CD19- [%]
CD8 SP/ CD5+CD19- [%]
DN/ CD5+CD19- [%]
DP/ CD5+CD19- [%]
T cell selection (E17.5)
[%] of CD5+ cells
DP
DN
CD4+
CD8+
WT
KO
ns
B
E18.5
Fam83h wt/wt
Fam83h -/-
range of intensity
Total volume [mm]
Total thymus space [mm]
Relative thymus volume [%]
C
DAPI + KRT5 + KRT8
Cortex
Medulla
100 µm
10 µm
E17.5
Area [µm2]
Area [%]
cTEC
mTEC

**Supporting Information Figure 5 | Immune cell development is unaffected in the distinct developmental stages in FL and thymus at embryonic timepoints E17.5 and E18.5.**

(A) Analysis of fetal hematopoiesis. Frequency of leukocyte subsets in the FL (upper panel, gating strategy shown in **Figure S9**) and thymocyte subsets in the thymus at E17.5 (gating strategy shown in **Figure S12;** DN (green) - double negative, DP (gray) - double positive, $CD4^+$ SP (blue, single positive), and $CD8^+$ SP (magenta)). Symbols represent individual embryos, and bars indicate mean ± SEM using an unpaired *t-test,* p-values: ns (P > 0.05). Sample sizes: *n*=8, 7 in FL; *n*=8, 7 in thymus for *Fam83h*$^{wt/wt}$ and *Fam83h*$^{-/-}$ mice, respectively.

(B) E18.5 embryos were imaged using µCT (left panel), and their volume, as well as thymic volumes, were calculated (graphs). At E18.5, *Fam83h*$^{-/-}$ embryos were not significantly smaller than littermates, and their thymus volumes were comparable. Symbols represent individual embryos, and bars indicate mean ± SEM using an unpaired *t-test,* p-values: ns (P > 0.05), *n*=4 for *Fam83h*$^{wt/wt}$ and *Fam83h*$^{-/-}$ mice.

(C) Cryosections from E17.5 embryonic stage in *Fam83h*$^{wt/wt}$ *and Fam83h*$^{-/-}$ thymi were stained for keratin-8 (cTEC, red) and keratin-5 (mTEC, green), revealing distinct cortical and medullary regions. DAPI staining marks nuclei, allowing differentiation between cortex and medulla. Representative images from 2 independent experiments are shown. Imaged with SDCM Nikon CSU-W1 Scale bars: 100 µm and 10 µm. Total thymic and cortical area was quantified from sections. Bars represent mean ± SD, n=4/3 for *Fam83h*$^{wt/wt}$ and *Fam83h*$^{-/-}$ embryos, respectively, unpaired *t-test.* p-value: ns (P > 0.05).

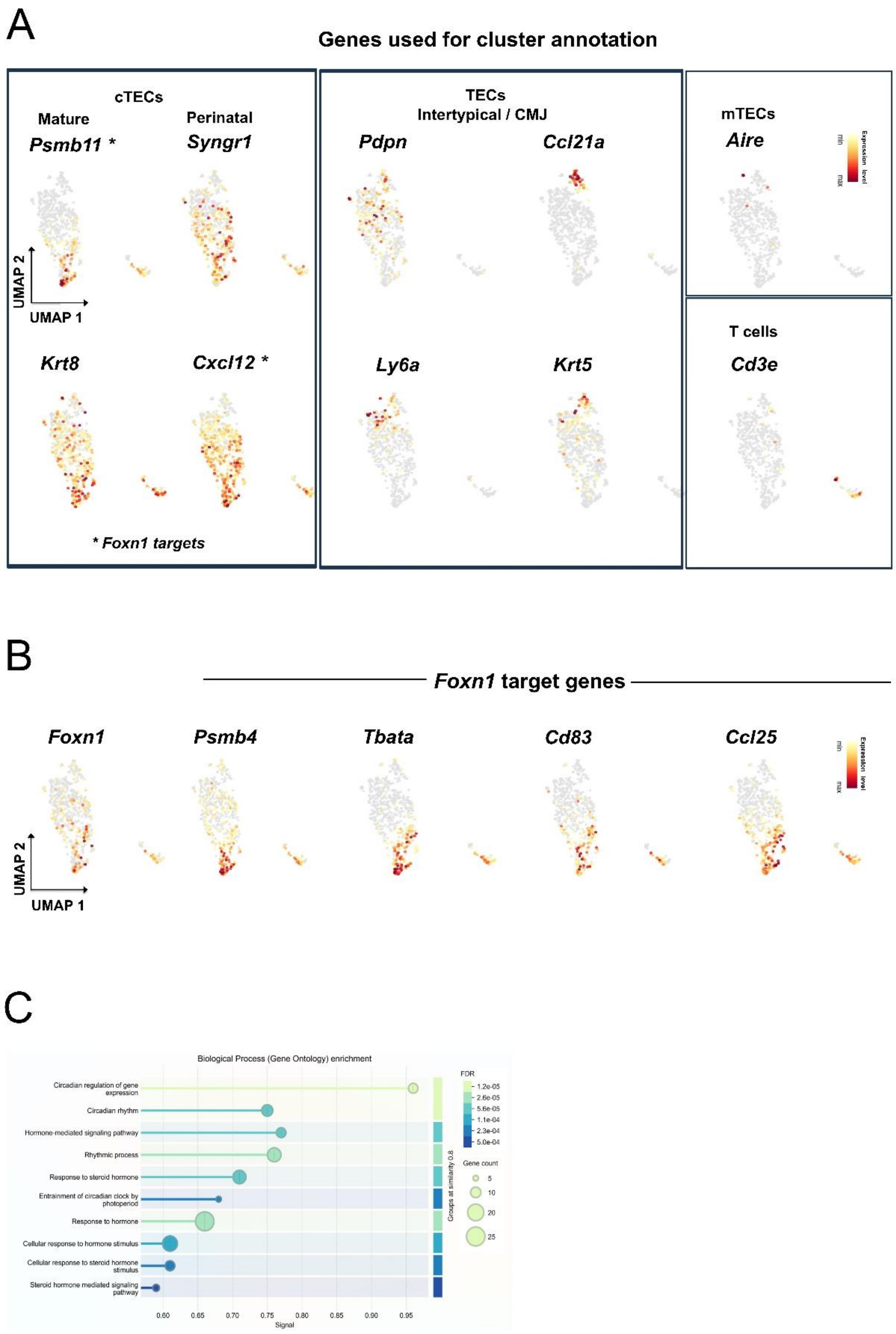
A
Genes used for cluster annotation
cTECs
Mature
Perinatal
Psmb11 *
Syngr1
UMAP 2
UMAP 1
Krt8
Cxcl12 *
* Foxn1 targets
TECs
Intertypical / CMJ
Pdpn
Ccl21a
Ly6a
Krt5
mTECs
Aire
Expression level
min
max
T cells
Cd3e
B
Foxn1 target genes
Foxn1
Psmb4
Tbata
Cd83
Ccl25
UMAP 2
UMAP 1
Expression level
min
max
C
Biological Process (Gene Ontology) enrichment
Circadian regulation of gene expression
Circadian rhythm
Hormone-mediated signaling pathway
Rhythmic process
Response to steroid hormone
Entrainment of circadian clock by photoperiod
Response to hormone
Cellular response to hormone stimulus
Cellular response to steroid hormone stimulus
Steroid hormone mediated signaling pathway
0.60
0.65
0.70
0.75
0.80
0.85
0.90
0.95
Signal
Groups at similarity 0.8
FDR
1.2e-05
2.6e-05
5.6e-05
1.1e-04
2.3e-04
5.0e-04
Gene count
5
10
20
25

**Supporting Information Figure 6 | Expression profiling of juvenile cTEC heterogeneity and *Foxn1* regulatory networks by scRNA-Seq.**

(A) Expression of selected marker genes used for cluster annotation according to (Baran-Gale, Morgan et al. 2020)

(B) Expression of *Foxn1* and *Foxn1*-regulated genes (*Psmb4, Tbata, Cd83, Ccl25*) is restricted to the most mature cTEC cluster.

(C) Identification of biological processes enriched in *Fam83h*$^{-/-}$ mice using selected genes obtained from Smart-Seq analysis (STRING database (https://string-db.org/)).

*Baran-Gale, J., M. D. Morgan, S. Maio, F. Dhalla, I. Calvo-Asensio, M. E. Deadman, A. E. Handel, A. Maynard, S. Chen, F. Green, R. V. Sit, N. F. Neff, S. Darmanis, W. Tan, A. P. May, J. C. Marioni, C. P. Ponting and G. A. Holländer (2020). "Ageing compromises mouse thymus function and remodels epithelial cell differentiation." Elife* ***9****.*

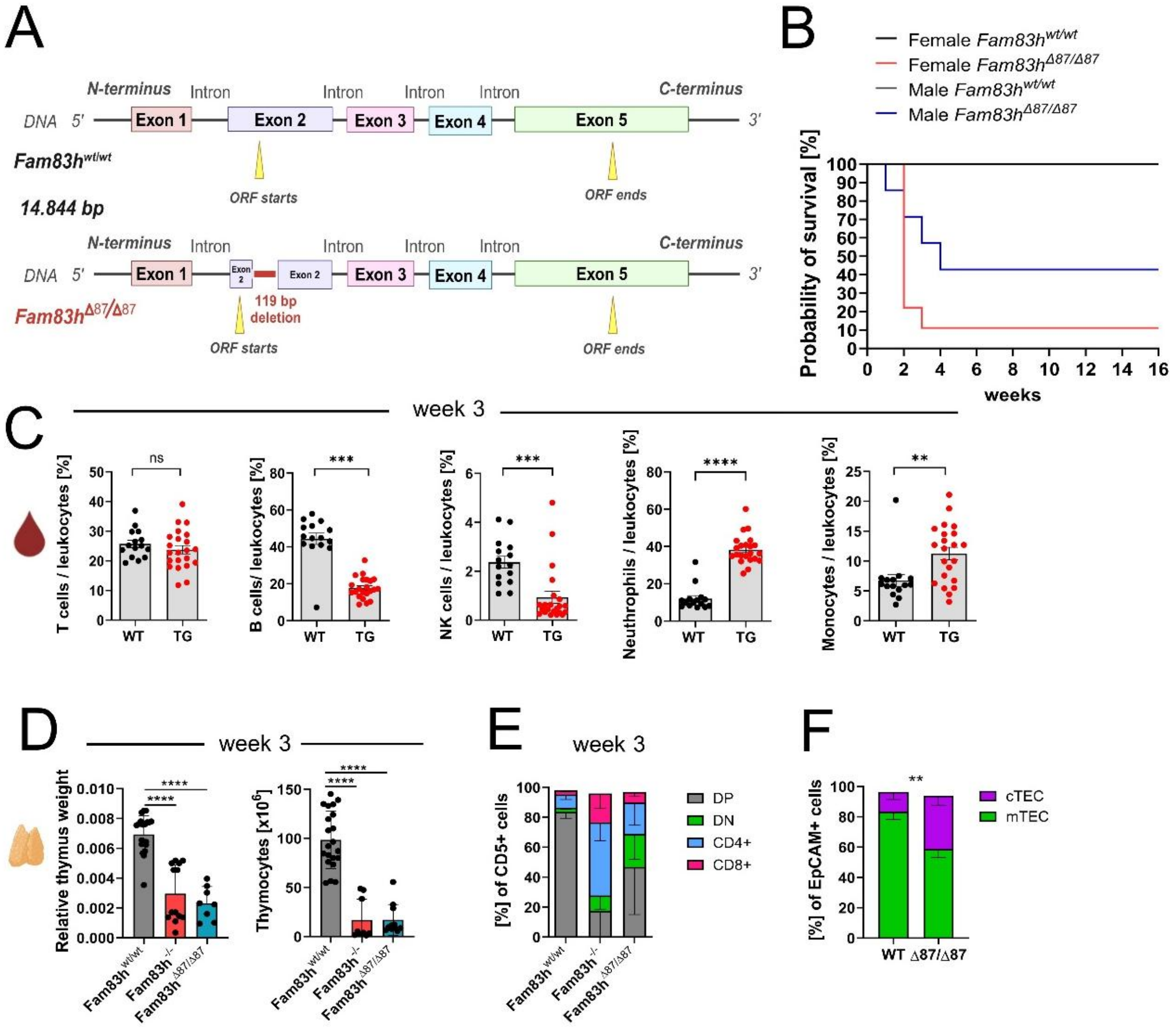

**Supporting Information Figure 7 | The truncated form of Fam83h (*Fam83h*$^{\Delta87/\Delta87}$) and *Fam83h*$^{-/-}$ exhibit a similar impact on hematopoiesis and T cell development.**

(A) Generation of *Fam83h*$^{\Delta87/\Delta87}$ mice – schematic representation showing a 119 bp deletion in exon 2 (created in BioRender).

(B) The survival curve shown represents the probability of survival over 16 weeks in male and female mice with different *Fam83h* genotypes: *Fam83h*$^{wt/wt}$, *Fam83h*$^{\Delta87/\Delta87-}$.

(C) Leukocyte percentage in peripheral blood was measured by FCM at week 3 of age. The full gating strategy is shown in **Figure S8.** The data represents 11 independent experiments. Bars indicate mean ± SEM, two-tailed unpaired *t-test*, p-values: ns (P > 0.05), ***P ≤ 0.001, ****P ≤ 0.0001. Sample sizes: *n*=15 for *Fam83h*$^{wt/wt}$ and n=22 for *Fam83h*$^{\Delta87/\Delta87}$ mice, respectively.

(D) Relative thymus size (left) and thymocyte counts (right) in *Fam83h*$^{wt/wt}$, *Fam83h*$^{-/-}$, and *Fam83h*$^{\Delta87/\Delta87}$ mice at week 3. The data represents 3 independent experiments. Bars indicate mean ± SD using one-way ANOVA, p-values: ****P ≤ 0.0001. Sample sizes: *n*=21/21, *n*=13/9 and *n*=8/11 for *Fam83h*$^{wt/wt}$, *Fam83h*$^{-/-}$ and *Fam83h*$^{\Delta87/\Delta87}$ mice, respectively.

(E) FCM analysis of the main thymocyte subsets. The full gating strategy is shown in **Figure S12**: DN (green), DP (gray), CD4$^{+}$ SP (blue), and CD8$^{+}$ SP (magenta) in *Fam83h*$^{wt/wt}$, *Fam83h*$^{-/-}$, and *Fam83h*$^{\Delta87/\Delta87}$ at week 3 of age. Bars indicate mean ± SD. Data are based on 3 independent experiments. Sample sizes: *n*=14, *n*=6 and *n*=10 for *Fam83h*$^{wt/wt}$, *Fam83h*$^{-/-}$, and *Fam83h*$^{\Delta87/\Delta87}$ mice, respectively.

(F) Percentage of cTECs and mTECs from EpCAM+ cells was determined by FCM. The gating strategy is shown in **Figure S17**. Bars represent mean ± SD, n=4/3 for *Fam83h*$^{wt/wt}$ and *Fam83h*$^{\Delta87/\Delta87}$ thymi respectively unpaired *t-test,* p-values: **P ≤ 0.01.

### Figure S8

#### Blood panel

***Fam83h*** $^{wt/wt}$

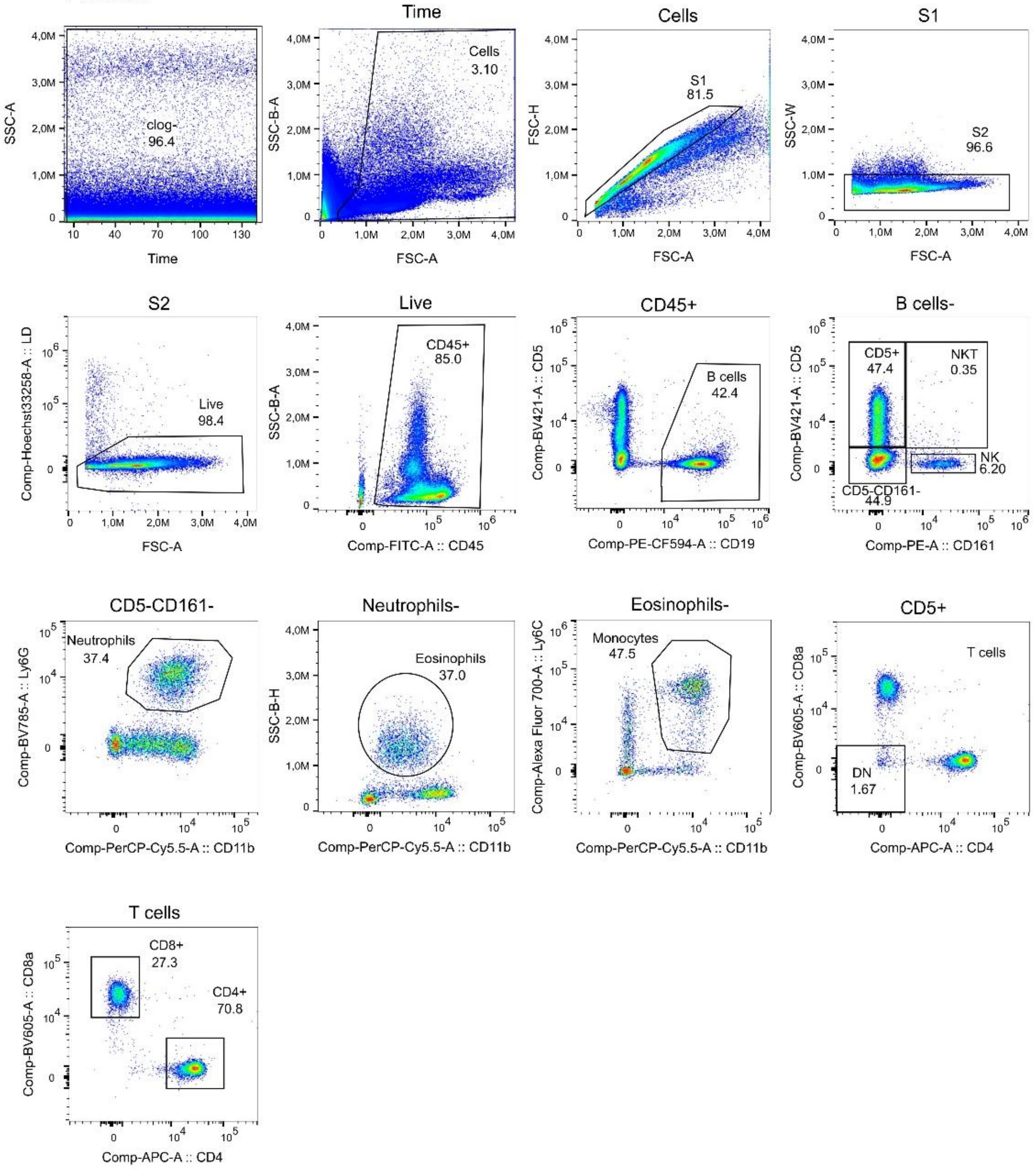

#### Figure S9
#### BM progenitor panel
***Fam83h*$^{wt/wt}$**

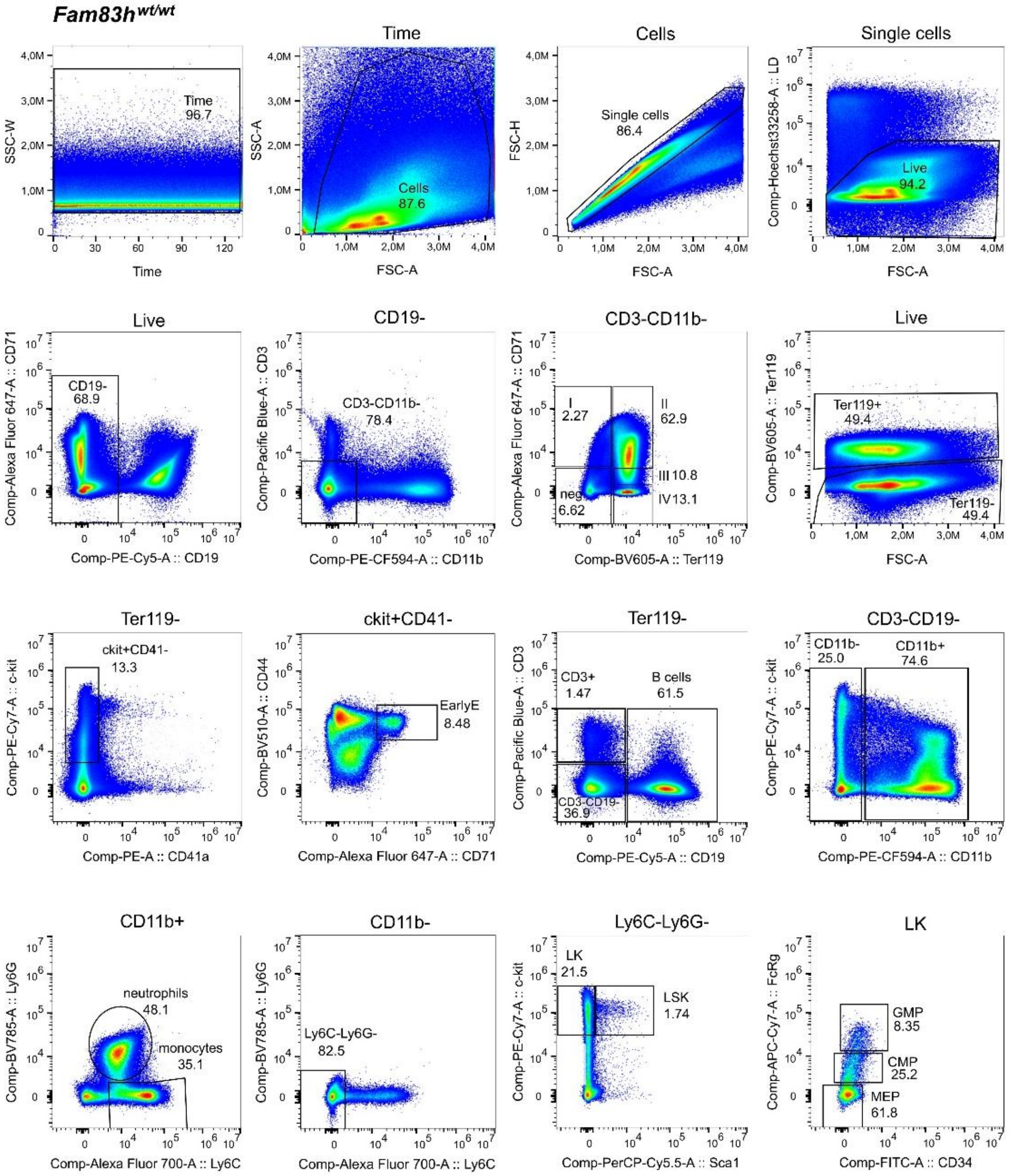

#### Figure S10

##### BM-B&NK progenitor panel

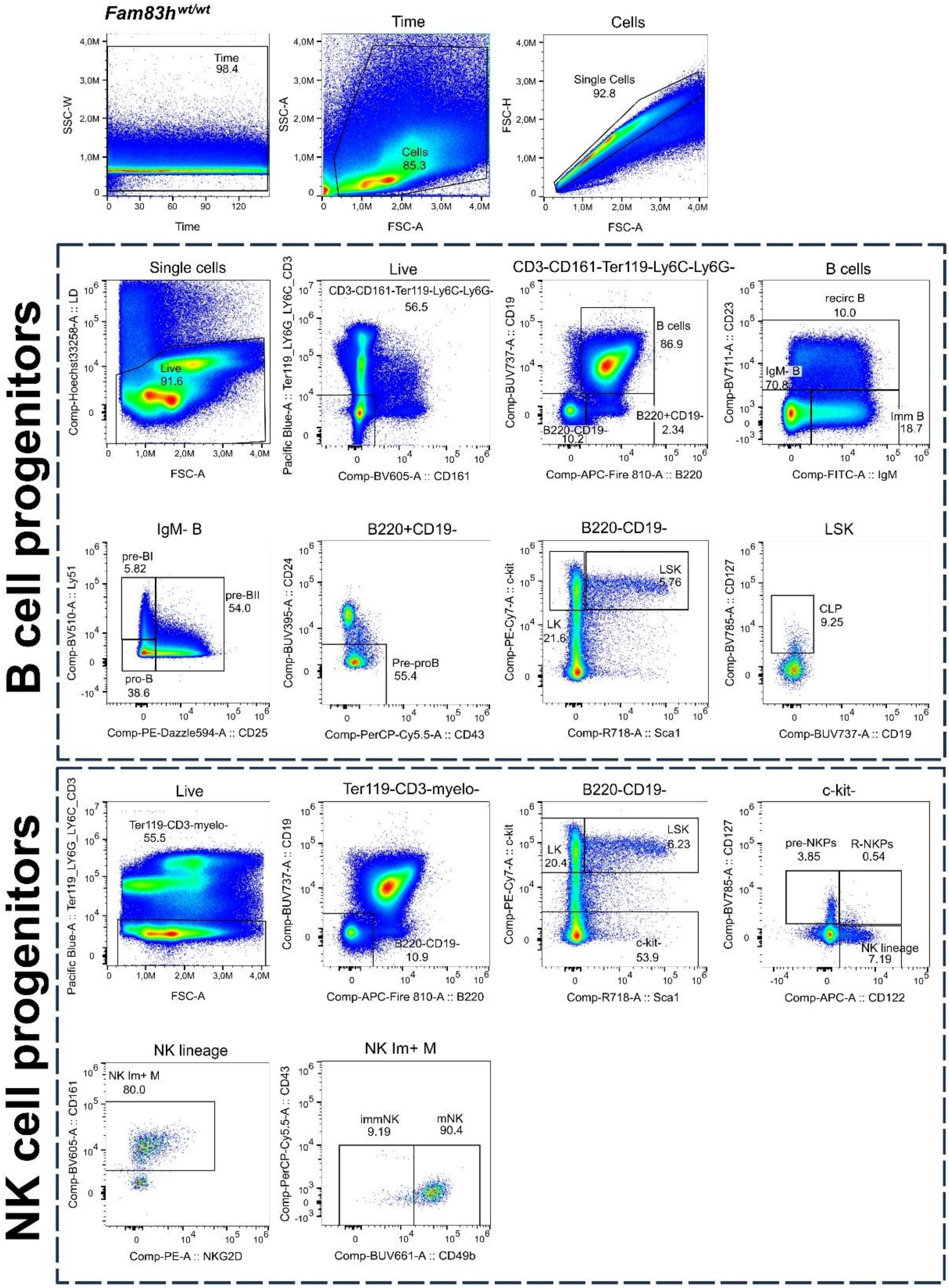

#### Figure S11

#### HPSC panel

***Fam83h**$^{wt/wt}$*

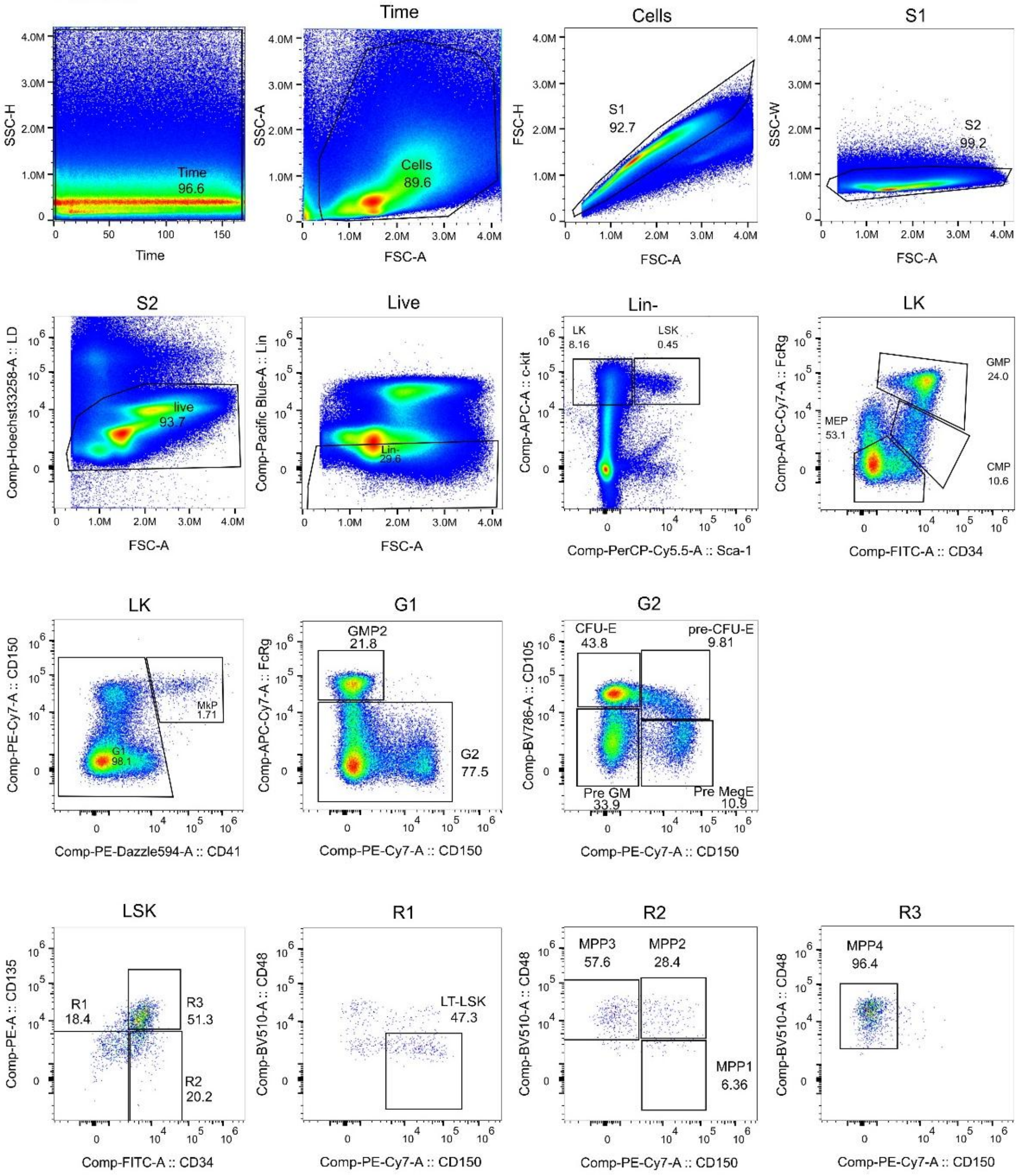

**Figure S12**

**Thymocyte panel**

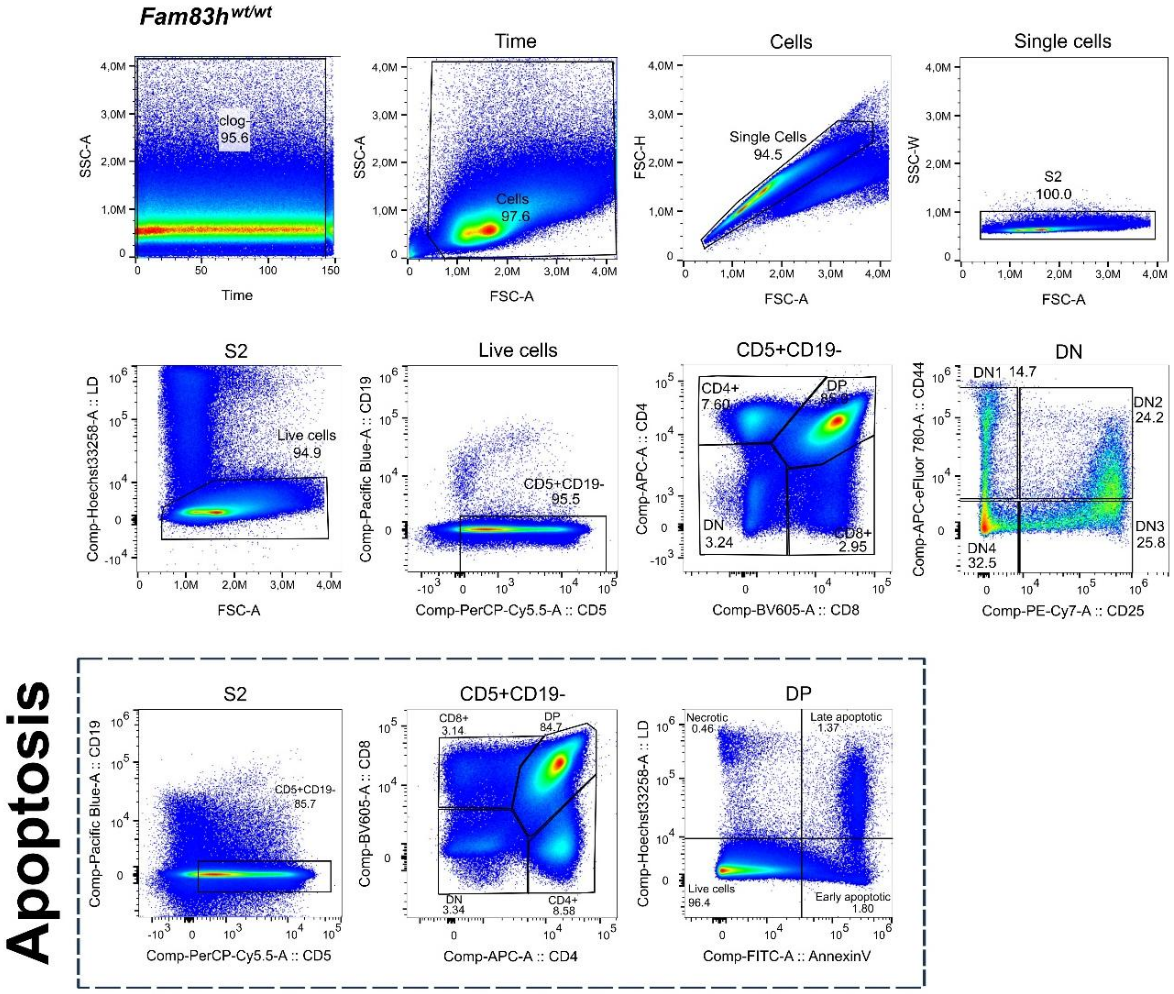

### Figure S13

#### Thymocyte-Ki67 panel

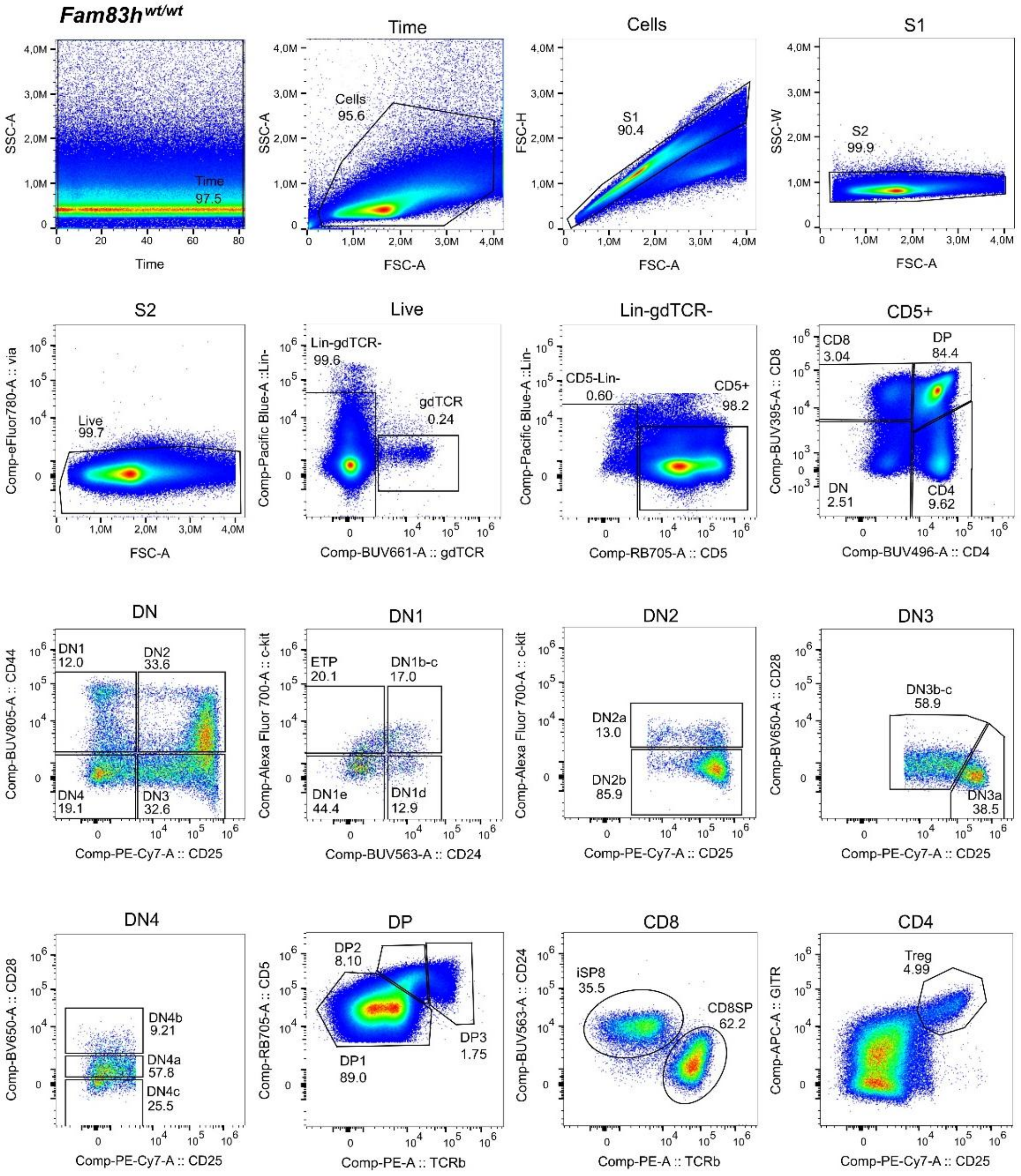


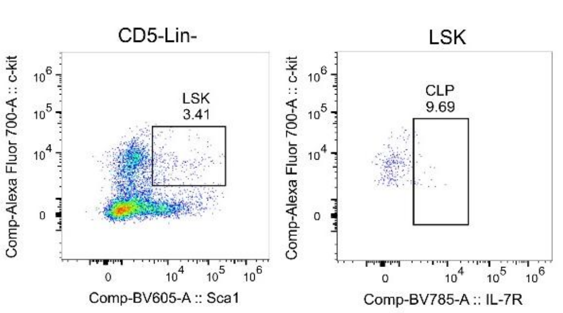


**DN/Ki67**

| | Sample Name | Subset Name | Count |
|---|---|---|---|
| | wt_F1497.fcs | DN3 | 10398 |
| | wt_F1496.fcs | DN3 | 8453 |
| | wt_F1485.fcs | DN3 | 6748 |
| | wt_F1484.fcs | DN3 | 10107 |
| | KO_F1504.fcs | DN3 | 2298 |
| | KO_F1502.fcs | DN3 | 13929 |
| | KO_F1487.fcs | DN3 | 8804 |
| | KO_F1486.fcs | DN3 | 916 |
| | iso_wt_F1484.fcs | DN3 | 7778 |

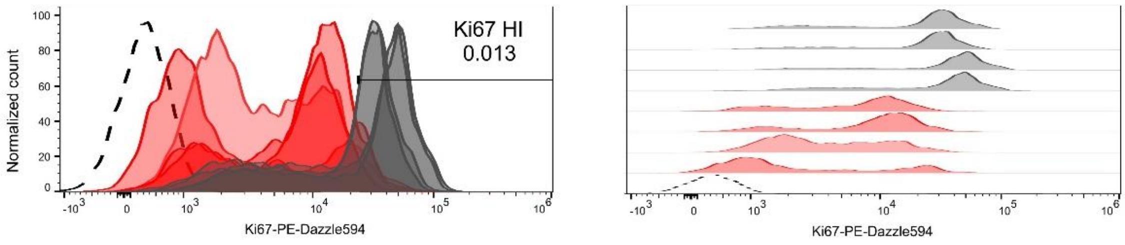


Lin: Ter119,CD11b,CD19,CD161

#### Figure S14

#### Chimera - thymocyte panel

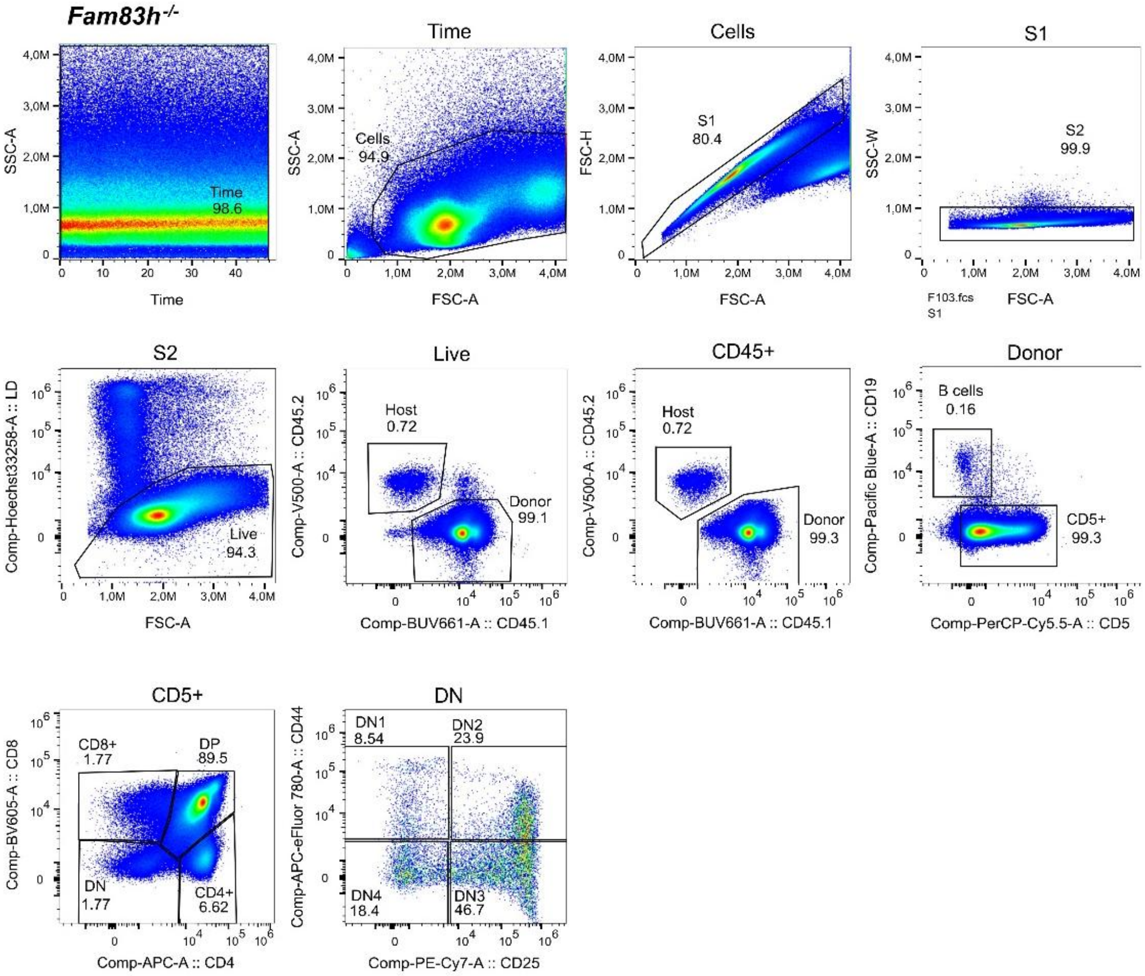

#### Figure S15

##### Chimera - Blood panel

***Fam83h$^{-/-}$***

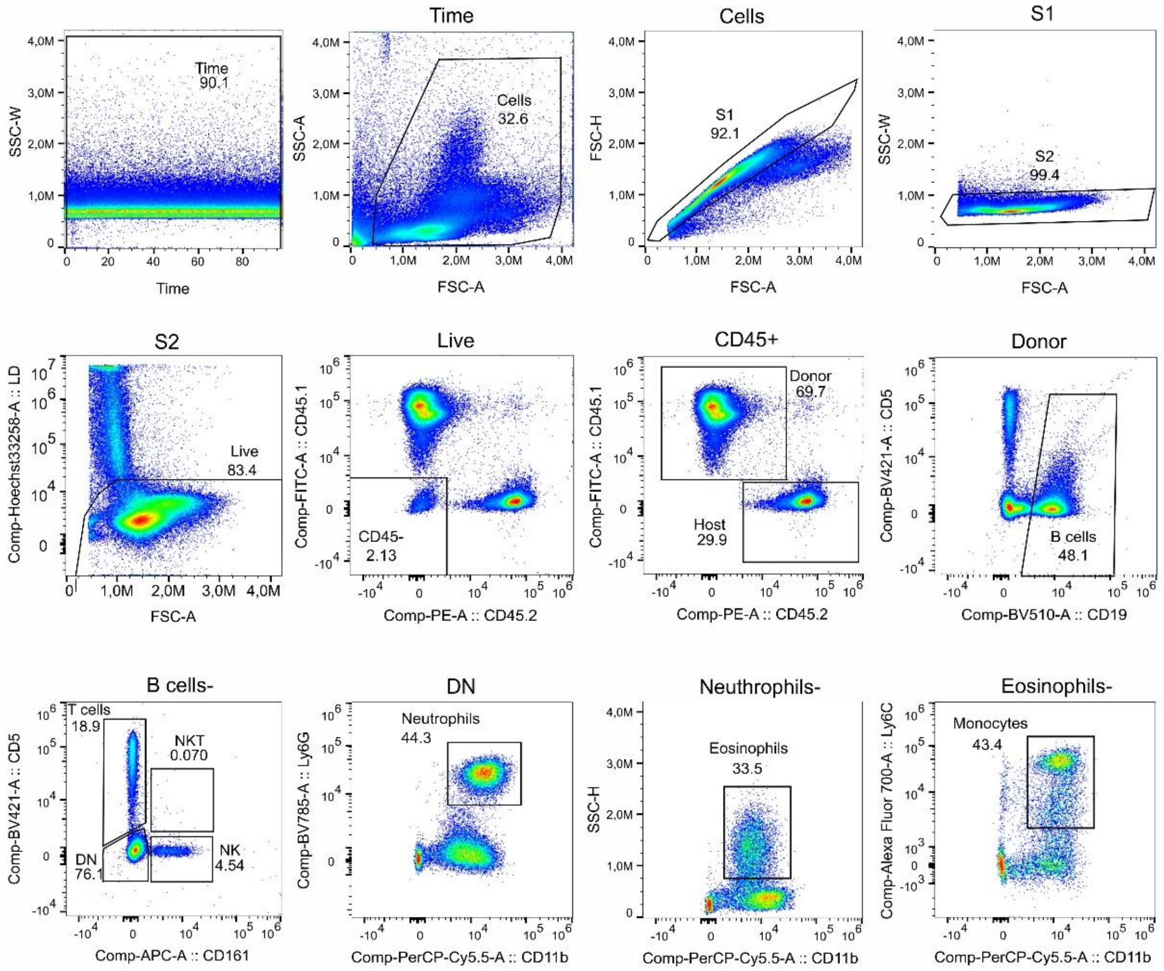

#### Figure S16

##### Chimera - B&NK panel

*Fam83h*$^{-/-}$

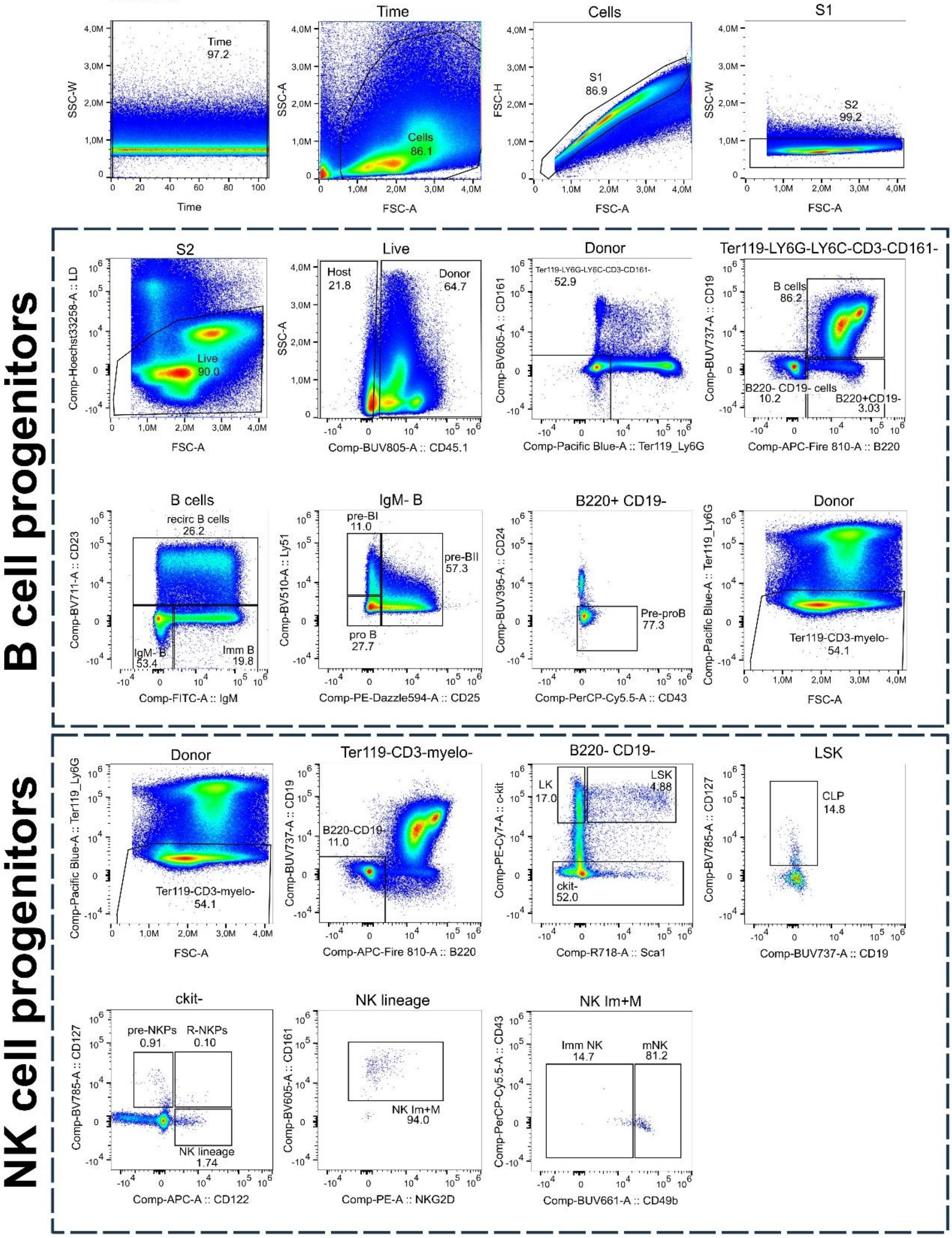

#### Figure S17

##### Thymus-TEC-sort panel

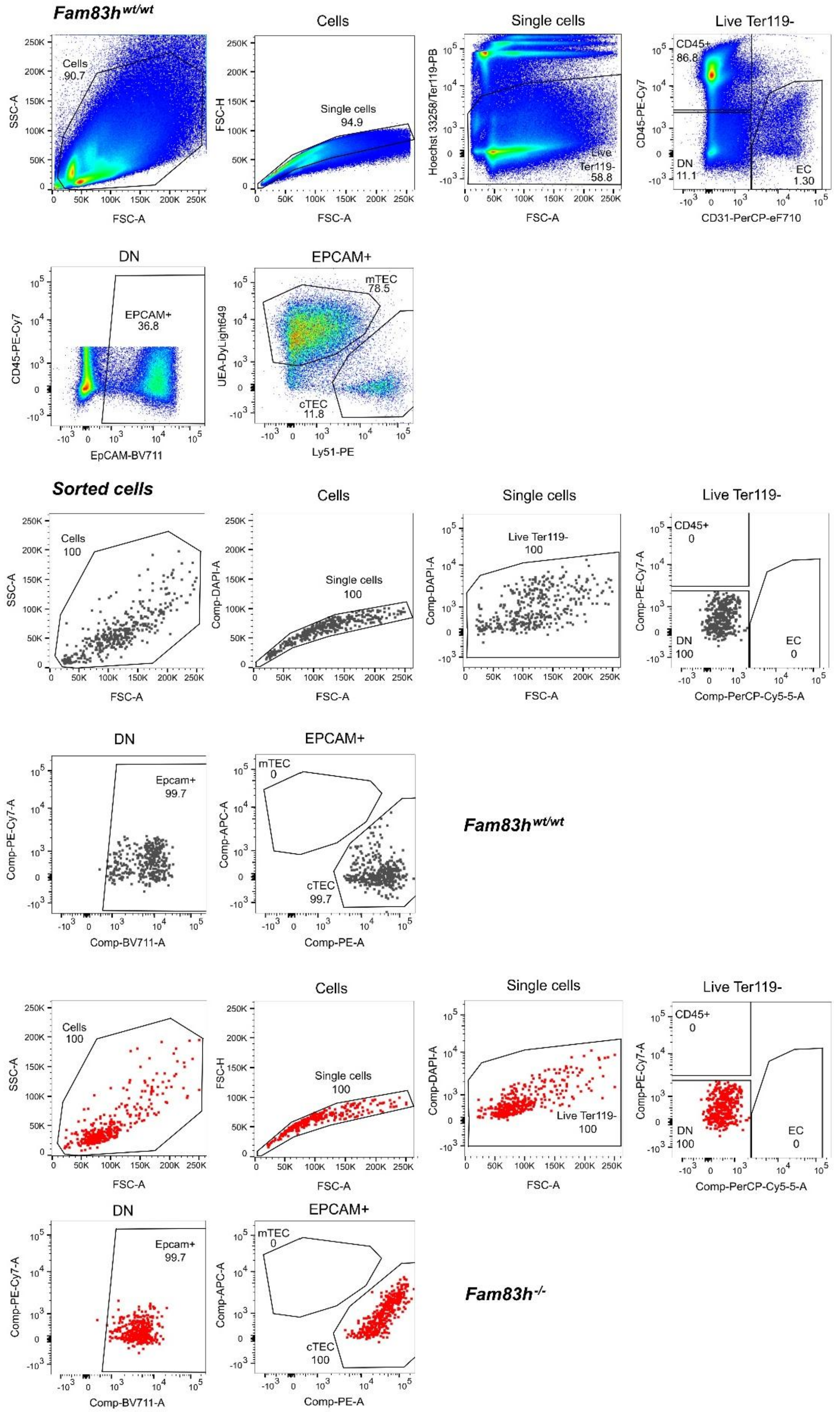

#### Figure S18

#### TEC-Ki67 panel

***Fam83h^wt/wt***

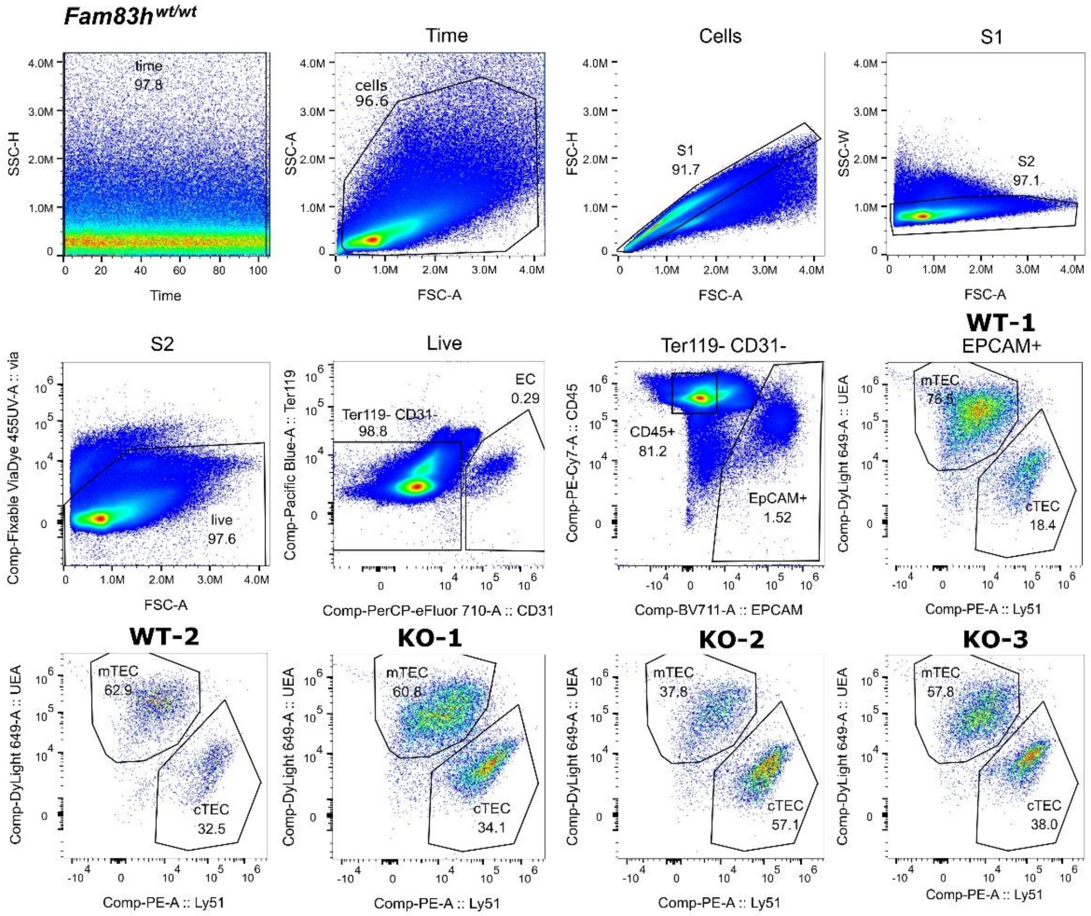


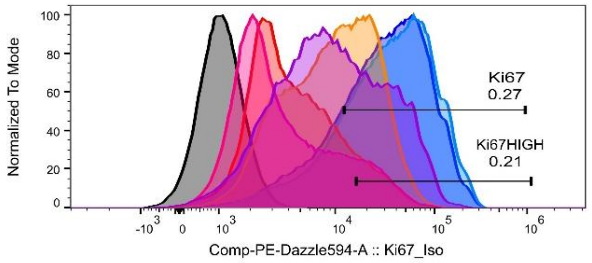


| | Sample Name | Subset Name | Count |
|---|---|---|---|
| | F1466_KO.fcs | CD45+ | 29643 |
| | F1454_KO.fcs | CD45+ | 326916 |
| | F1451_KO.fcs | CD45+ | 260665 |
| | A_KO.fcs | CD45+ | 55768 |
| | F1467_WT.fcs | CD45+ | 8.34E5 |
| | F1464_WT.fcs | CD45+ | 199482 |
| | F1464_WT_Isotype.fcs | CD45+ | 150201 |

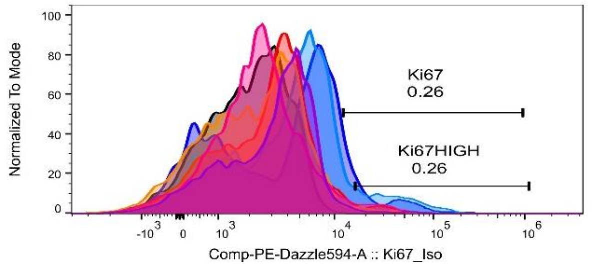


| | Sample Name | Subset Name | Count |
|---|---|---|---|
| | F1466_KO.fcs | cTEC | 3732 |
| | F1454_KO.fcs | cTEC | 5398 |
| | F1451_KO.fcs | cTEC | 4738 |
| | A_KO.fcs | cTEC | 5045 |
| | F1467_WT.fcs | cTEC | 3179 |
| | F1464_WT.fcs | cTEC | 1672 |
| | F1464_WT_Isotype.fcs | cTEC | 1150 |

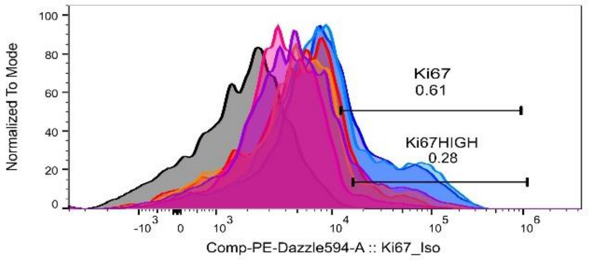


| | Sample Name | Subset Name | Count |
|---|---|---|---|
| | F1466_KO.fcs | mTEC | 5535 |
| | F1454_KO.fcs | mTEC | 9540 |
| | F1451_KO.fcs | mTEC | 3102 |
| | A_KO.fcs | mTEC | 2729 |
| | F1467_WT.fcs | mTEC | 12451 |
| | F1464_WT.fcs | mTEC | 3104 |
| | F1464_WT_Isotype.fcs | mTEC | 2143 |

### Supplementary Tables

**Gel electrophoresis**: 12uL on 2% agarose gel, 90V, 45min.

**DNA isolation:** Phenol/Chloroform (for E17.5), and Quick Extract DNA extraction solution

(#1) PCR product (#1) F1xR1 for wt 741bp, F1xR1 for mutated 622bp.

(#2) PCR product (#2) F1x R1 for wt 6061 (no deletion), 1134bp with deletion.

| | **Forward primer** | **Reverse primer** |
|---|---|---|
| ***Fam83h (#1)*** | 5'-ACTGGCCCATGTCTTACCAG -3' | 5'-CTCCCACACCCTCCTATTCA-3' |
| ***Fam83h-GD (#2)*** | 5'-CAGATGCAGGTGTGGACTGA -3' | 5'-GAACGTCTGTCCCGATTCCT-3' |

**Supplementary Table 1 | Primer sequences and PCR conditions used for genotyping *Fam83h* alleles.**

| | **Forward** | **Reverse** |
|---|---|---|
| ***Fam83h*** | GTGGGCCAGAAGCCTACAAC | GCTCACGTGTTCCAGTTCCT |
| ***Casc3* (RF)** | TTCGAGGTGTGCCTAACCA | GCTTAGCTCGACCACTCTGG |
| ***Foxn1*** | CCAGCCAGGAACACAACCAA | GGCAGGCTGAGAAGAACAGT |
| ***Ccl2*** | GCTACAAGAGGATCACCAGCAG | GTCTGGACCCATTCCTTCTTGG |
| ***Per1*** | GAAACCTCTGGCTGTTCCTACC | AGGCTGAAGAGGCAGTGTAGGA |
| ***Txnip*** | GTTGCGTAGACTACTGGGTGAAG | CTCCTTTTTGGCAGACACTGGTG |
| ***Ccl21a*** | CCCTACAGTATTGTCCGAGGC | TCAGGCTTAGAGTGCTTCCG |
| ***Psmb11*** | CCACCCACCGTGATGCTTAT | GCTCCCTATACAGCACGCAA |
| ***Ajap1*** | AAGGTCTGAGGCTGGGTTCC | GATGGGAAGTCGACCGCAAG |

**Supplementary Table 2 | List of Primers Used for RT-qPCR**

| Specificity/ viability | Clone | Host | Company | Catalog # |
|---|---|---|---|---|
| DAPI | n/a | n/a | Sigma-Aldrich | 10236276001 |
| Anti-KRT8 (Keratin 8) | TROMA-1 | Rat | Sigma-Aldrich | MABT329 |
| Anti-KRT5 (Keratin 5) | 3P3T9 | Rabbit | ThermoFisher | MA5-35103 |
| Anti-E-cadherin | DECMA-1 | Rat | ThermoFisher | 13-1900 |
| Anti-ZO-1 | Polyclonal | Rabbit | ThermoFisher | 40-2200 |
| Anti-VCAM-1 | SA05-04 | Rabbit | ThermoFisher | MA5-31965 |
| Anti-Rat IgG1 κ Isotype Control | eBRG1 | Rat | ThermoFisher | 14-4301-82 |
| Anti-Rabbit IgG Isotype Control | Polyclonal | Rabbit | ThermoFisher | 31235 |

**Supplementary Table 3 | List of Antibodies Used in Histological Staining (H&E, IF)**

| Specificity/ viability | Fluorochrome | Clone | Company | Catalog # |
|---|---|---|---|---|
| Fc Block | | 2.4G2 | BD | 553142 |
| Ly6C | AF700 | AL-21 | BD | 561237 |
| CD5 | BV421 | 53-7.3 | BD | 562739 |
| CD44 | BV510 | IM7 | Biolegend | 103043 |
| Ly6G | BV785 | 1A8 | Biolegend | 127645 |
| GITR | BV711 | DTA-1 | BD | 563390 |
| CD25 | PE-Cy7 | PC61 | BD | 552880 |
| CD8a | BV605 | 53-6.7 | Biolegend | 100744 |
| CD45 | FITC | 30-F11 | Biolegend | 103108 |
| CD11b | PerCPCy5.5 | M1/70 | Biolegend | 101228 |
| CD161 | PE | PK136 | SONY | 1143540 |
| CD19 | PE-CF594 | 1D3 | BD | 562291 |
| CD4 | APC | RM4-5 | SONY | 1102575 |
| CD62L | APC-Cy7 | MEL-14 | BD | 560514 |

| | | | | |
|---|---|---|---|---|
| TCRbeta | PE | H57-597 | Thermo Scientific | 12-5961-81 |
| CD19 | PB | 6D5 | Biolegend | 115523 |
| CD44 | APC-eF780 | IM7 | Thermo Scientific | 47-0441-80 |
| Annexin V | FITC | B266195 | Biolegend | 640906 |
| CD5 | PerCPCy5.5 | 53-7.3 | SONY | 1103115 |
| CD3 | PB | 17A2 | Biolegend | 100214 |
| FcRg (CD16/32) | APC-Cy7 | 93 | Biolegend | 101328 |
| c-kit | PC7 | ACK2 | SONY | 1275555 |
| Ter119 | BV605 | TER-119 | Biolegend | 116239 |
| CD34 | FITC | RAM34 | Invitrogen | 11034182 |
| Sca1 | PerCPCy5.5 | D7 | SONY | 1140620 |
| CD19 | PE-Cy5 | 6D5 | Biolegend | 115509 |
| CD11b | PE-CF594 | M1/70 | BD | 562287 |
| CD41a | PE | MWReg30 | Thermo Scientific | 12-0411-81 |
| CD71 | AF647 | RI7217 | prepared in-house | n/a |
| CD24 | BUV395 | M1/69 | BD | 744471 |
| CD49b | BUV661 | HMa2 | BD | 741523 |
| CD19 | BUV737 | 1D3 | BD | 612782 |
| Ter119 | PB | TER-119 | Biolegend | 116232 |
| Ly6G, Ly6C (Gr-1) | PB | RB6-8C5 | Biolegend | 108430 |
| BP1 (Ly51) | BV510 | BP-1 | BD | 745061 |
| CD161 | BV605 | PK136 | BD | 563220 |
| CD23 | BV711 | B3B4 | BD | 563987 |
| IL-7Ra? (CD127) | BV785 | A7R34 | Biolegend | 135037 |
| IgM | FITC | RMM-1 | Biolegend | 406506 |
| CD43 | PerCPCy5.5 | 1B11 | Biolegend | 121223 |
| NKG2D | PE | CX5 | BD | 558403 |
| CD25 | PEDazz594 | PC61 | Biolegend | 102048 |
| CD122 | APC | TM-β1 | Biolegend | 123213 |
| Sca1 | R718 | D7 | BD | 567733 |

| | | | | |
|---|---|---|---|---|
| B220 | APC-Fire810 | RA3-6B2 | Biolegend | 103277 |
| Fixable Viability Dye eFluor™ | UV455 | | Thermo Scientific | 65-0868-18 |
| EpCAM | BV711 | G8.8 | Biolegend | 118233 |
| CD31 | PerCP-ef710 | 390 | eBioscience | 46-0311-80 |
| Ly6D | FITC | 49-H4 | BD | 561148 |
| Ly51 | PE | 6C3 | Biolegend | 108307 |
| CD45 | PC7 | 30-F11 | Biolegend | 103114 |
| UEA | DyLight649 | n/a | Baria | DL-1068-1 |
| MHCII | APC-Cy7 | M5/114.15.2 | Biolegend | 107628 |
| Ki67 | PEDazzle594 | 11F6 | Biolegend | 151219 |
| IgG1 isotype | PEDazzle594 | RTK2071 | Biolegend | 400445 |
| Hoechst 33258 | n/a | n/a | Sigma-Aldrich | 94403 |
| CD45 | PB | 30-F11 | Biolegend | 103126 |
| CD105 | BV786 | MJ7/18 | BD | 564746 |
| CD31 | FITC | 390 | Biolegend | 102435 |
| Sca1 | PercpCy5.5 | D7 | eBioscience | 45-5981-80 |
| CD51 | PE | RMW-7 | Thermo Scientific | 12-0512-81 |
| CD71 | PE-Dazz594 | RI7217 | SONY | 1169085 |
| Vcam | PC7 | 429 | Biolegend | 105719 |
| c-kit | APC | 2B8 | Biolegend | 105812 |

**Supplementary Table 4 | List of Antibodies Used for Flow Cytometry Analysis**

| **Specificity/ viability** | **Clone** | **Host** | **Company** | **Catalog #** |
|---|---|---|---|---|
| Anti-FAM83H | Polyclonal | Rabbit | Novus | NBP2-32219 |
| Anti-CK1a | Polyclonal | Rabbit | Proteintech | 55192-1-AP |

**Supplementary Table 5 | List of Antibodies Used for Co-IP**
